# Predicting Supramolecular Self-Assembly of Peptide Structures with AlphaFold3

**DOI:** 10.64898/2026.04.28.720402

**Authors:** Claire Sklar, Jessie Huh, Sabrina Chen, Jeffrey J. Gray

**Affiliations:** Department of Chemical and Biomolecular Engineering, Johns Hopkins University, Baltimore, MD, 21218, United States

## Abstract

Self-assembled peptide-based nanostructures have diverse applications in the pharmaceutical and materials fields, but accurately predicting their self-assembly behavior without time-intensive organic synthesis and characterization remains a significant challenge. Here, we assess the effectiveness of AlphaFold3 (AF3), a deep learning model for protein structure prediction, in modeling peptide-based nanostructures and the interactions driving supramolecular self-assembly. We designed amphiphilic peptides composed of alternating hydrophobic residues (valine, leucine, isoleucine, phenylalanine) and hydrophilic residues (glutamic acid), varying both sequence length and residue order. Using AF3’s multimer mode, we modeled assemblies with copy numbers ranging from 10 to 1000, generating diverse morphologies such as micelles and nanotubes. We qualitatively analyzed hydrophobic regions, secondary structures, and intermolecular interactions, while also calculating radii of gyration, packing scores, and aspect ratios using PyRosetta. Our results indicate that AF3 predicts morphologies consistent with hydrophobic driving forces and steric constraints. Increased hydrophobicity correlates with smaller radii of gyration, while higher copy numbers correspond to smaller aspect ratios (more compact structures). Longer hydrophobic segments lead to disordered structures, whereas longer hydrophilic segments promote organization. While AF3 captures systemic trends consistent with biophysical principles, comparisons to literature reveal discrepancies driven by charge effects and secondary structure bias, including an overemphasis on helical propensity (e.g., alanine-rich sequences) and sensitivity to terminal charge repulsion. Additionally, since AF3 is predisposed to predict a single assembled entity rather than higher-order assemblies such as multiple micelles or fibers, finding the optimal copy number for the best prediction requires system-specific iteration. These limitations highlight the need for complementary approaches with controlled chemical potential and environmental conditions, though qualitative agreement with experimental trends in morphology and compactness supports AF3’s utility for initial structure generation. Our findings highlight AF3’s potential as a user-friendly first pass screening tool for peptide design, enabling rapid generation and prioritization of candidate assemblies for subsequent evaluation using coarse-grained or all-atom molecular simulations toward the efficient development of functional self-assembled peptide nanomaterials.

## Introduction

Self-assembled, peptide-based nanostructures have emerged as versatile materials for applications spanning biomaterials, electronics, sensing, catalysis, interfacial engineering, and pharmaceuticals. These include surface coatings with antifouling or selective adhesion characteristics,^1–6^ platforms for tissue engineering and regenerative medicine,^7–9^ conductive materials,^10–13^ biological and environmental sensors,^14–16^ delivery systems for food and pharmaceuticals,^17–24^ implantable and injectable biomaterials,^13,25^ and integrated MOFs or COFs for separation and catalysis.^26–28^ Modifying the peptide design and assembly conditions allows engineers to direct the self-assembly of the structures into diverse morphologies (nanofibers,^29–31^ nanotubes,^32–34^ vesicles,^32,35^ nanofilaments,^36^ micelles,^37^ and nano-doughnuts^38^) and subsequently tune their functionality. This structural diversity underlies the broad utility of peptide materials but also makes predicting and controlling self-assembly one of the central challenges in the field.

The rational design of self-assembling peptides remains difficult because assembly emerges from a complex interplay among amino acid sequence, intermolecular interactions, and environmental conditions. Small changes in sequence,^31,39–54^ concentration,^39,55–57^ pH,^17,39,58–60^ or solvent^55,56,61–63^ can produce dramatically different supramolecular morphologies. Thus, experimental discovery is highly iterative and is often limited to the experimental time scale of synthesis, purification, and characterization. Computational approaches have become increasingly attractive for accelerating peptide design, yet accurately predicting self-assembly remains an unresolved challenge. Approaches combining coarse-grained (e.g., MARTINI^64^, OPEP^65^) and all-atom simulations (e.g., CHARMM^66^, GROMACS^67^), often standardizing concentration and environment polarity (e.g., water with NaCl), have been widely employed to link self-assembly to aggregation propensity.^68,69^ These approaches have reproduced important experimental trends, including the effects of terminal charge, the impact of sequence length, and the importance of charged residue placement.^53^ Despite these advances, current computational approaches remain limited in their ability to generate broadly transferable design rules. Coarse-grained models simplify residue interactions to access larger system sizes and longer timescales but sacrifice atomic detail needed to accurately capture electrostatic interactions, hydrogen bonding, and conformational dynamics. Conversely, all-atom approaches provide this molecular resolution but remain computationally expensive, limiting accessible system sizes, sampling times, and high-throughput exploration of peptide sequence space.^70^ As a result, many computational studies are necessarily performed under narrowly defined chemical and environmental conditions, making it difficult to generalize conclusions across diverse peptide chemistries, solvent environments, or target morphologies.

For example, Frederix et al. used the coarse-grained MARTINI force field to screen 8,000 tripeptide sequences and derive a hydrophobicity-based aggregation propensity score, identifying sequence preferences such as aromatic residues at positions 2 and 3, positively charged and hydrogen-bonding residues at the N-terminus, and negatively charged residues at the C-terminus.^53^ While these results demonstrated the utility of large-scale computational screening, the resulting design rules were derived under a standardized solvent environment and primarily reflected hydrophobicity-driven assembly. The authors noted opportunities to extend the framework by incorporating additional descriptors, including nanostructure morphology, peptide foldability, and specific intermolecular interactions, highlighting the need for more comprehensive and transferable predictive models. Similarly, Sarma et al. engineered a Monte Carlo coarse-grained pipeline to identify peptides that self-assemble into β-sheet fibrils.^71^ The simulations demonstrated computational control over β-strand organization within β-sheets, but directly relating the computational predictions to experimental CD and FTIR measurements remained challenging, illustrating the broader difficulty of connecting simulation outputs with experimentally observable assembly behavior. Collectively, these studies demonstrate that existing simulation frameworks can successfully interrogate specific peptide systems, but they remain limited in their ability to generate broadly transferable design rules and sequence-structure-property relationships. Developing computational models that balance molecular accuracy, computational efficiency, and generalizability across diverse peptide chemistries remains a major challenge for the rational design of self-assembling peptide materials.

Advances in AlphaFold3 (AF3) offer exciting opportunities for modeling peptide self-assembly. AF3 has shown high accuracy in predicting individual protein structures, even integrating information about molecular interactions beyond individual polypeptides and foreshadowing successful prediction of higher-order assembly. In multimer mode, AF3 models multiple copies of peptide sequences as interacting chains within a unified complex, allowing it to predict quaternary structures. Importantly, AF3 does not require user-defined docking constraints or prior information about assembly interfaces; instead, it infers plausible multimeric arrangements directly from sequence data. Furthermore, its user-friendly web interface allows anyone to generate predictions, enabling widespread adoption with minimal training.

The application of AF3 to peptide self-assembly remains unexplored and presents important challenges. AF3 was developed and trained primarily on folded proteins and biomolecular complexes, where it has demonstrated strong performance in predicting intermolecular interfaces and complex structures.^72^ However, supramolecular peptide assembly often involves larger numbers of interacting molecules, with assembly behavior governed by collective intermolecular interactions, peptide concentration, solvent environment, and thermodynamic equilibrium. These factors are not explicitly represented within AF3, and the model requires the user to specify the number of peptide chains included in each prediction. Therefore, the ability of AF3 to predict peptide nanostructures and the extent to which predicted assemblies correspond to experimental results remains unknown.

In this study, we used AlphaFold3 as the primary tool for structure generation, and we analyzed the outputs using complementary computational and visualization software. PyRosetta^73^ is a Python-based interface to the Rosetta, a molecular modeling suite. PyRosetta is used for further analysis of AF3-predicted assemblies, including evaluation of residue-level features, interchain distances, and assembly compactness. MDAnalysis,^74,75^ a computational tool for the analysis of molecular simulation data, is also used to calculate structural descriptors such as radius of gyration from the predicted assemblies. In support, PyMOL^76^ is used for molecular visualization of the peptide structures. Together, we use a workflow in which AlphaFold3 generates peptide assembly candidates, PyRosetta allows quantitative analyses of these outputs, and PyMOL supports visual inspection of the assemblies.

Here, we assess the utility of AF3 in predicting the interactions of peptide-based nanostructures, specifically in how the hydrophobic effect and other peptide-peptide interactions drive supramolecular self assembly. We take two complementary approaches. (1) We test a rational series of peptide designs featuring amino acids of increasing hydrophobicity (P, A, V, L, I, and F) complexed with hydrophilic glutamic acid (E), analyzed at different copy numbers. (2) We test specific sequences that have been characterized experimentally to directly compare the AF3 predictions with real data. We seek to probe AlphaFold3’s potential as a user-friendly design tool for structure generation in peptide design, aiding the development of functional self-assembled peptide nanomaterials.

### Experimental Section

Our workflow for this process was as follows: we submitted the input sequences to the AF3 server^72^ at varying copy numbers. Server access from April 2025 to December 2025 corresponded to AF3 version 3.0.1. Using the predicted structures, we visualized the outputs in PyMOL to identify hydrophobic regions, secondary structure, and intermolecular interactions. We performed quantitative structural analysis with PyRosetta to answer how generated structures conform to known principles of self-assembly (**Figure S1**). That is, we calculated trends in interface burial of hydrophobic regions, secondary structure formation (β-sheets and α-helices), intermolecular hydrogen bonding, and symmetric packing. We expanded this to expected principles and prior experimental studies on supramolecular growth. It is important to note that throughout this work, assembly behavior is analyzed as a function of peptide copy number rather than bulk concentration. Although increasing the number of peptide chains at a fixed volume would correspond to an increase in concentration, the effective volume represented by AlphaFold3 is not defined. Thus, a quantitative conversion between copy number and concentration can not be made.

### Sequence Design: Rationale

We evaluated AF3’s ability to predict self-assembly behavior of short amphiphilic peptides by designing a series of synthetic peptide sequences composed of distinct hydrophilic and hydrophobic segments. As our base case, we selected valine (V), which is hydrophobic and has a ß-branched side chain, and glutamic acid (E), which is hydrophilic and has a negatively charged side chain. We also investigated proline (P), alanine (A), leucine (L), isoleucine (I), and phenylalanine (F). We designed sequences that varied in length (e.g., 4 to 32 residues) and in the arrangement of hydrophobic and hydrophilic residues. These designs include block sequences such as VVVV…EEEE (V9E9) and alternating motifs such as VVVVEEEE…VVVVEEEE ((V4E4)4 ). We submitted each sequence to the AlphaFold3 server in varying quantities (10, 50, 100, 150, 200, 250, 300, 350, 500, and/or 1000 identical peptide chains) to approximate bulk self-assembly conditions. AlphaFold3 produces five predicted models for each input; for consistency, we chose the last structure predicted by AlphaFold3 for our analysis.

### Qualitative Analysis with PyMol Visualization

To visualize the predicted structural models, we used PyMOL versions 2.6 to 3.0.^76^ To assess spatial organization, we colored hydrophobic residues red and hydrophilic residues blue using the following command: color [color], resn [3-letter-code]. We used visualization to examine aggregation patterns, hydrophobic core alignment, and residue orientation. We then compared across different copy numbers and sequence designs to evaluate self-assembly tendencies and identify qualitative structural trends.

### Quantitative Analysis with PyRosetta and MDAnalysis

We analyzed the multimeric protein assemblies predicted by AF3 using PyRosetta^73^ and MDAnalysis.^74,75^ For each predicted structure, we identified individual monomer chains and processed them independently. We first calculated the radius of gyration of the full multimeric structure (**SI Eq. 1**) which provides a global measure of chain and assembly compactness. The radius of gyration provides a description of the overall structural expansion or compaction, with lower values indicating tighter packing and higher values indicating more extended conformations. Radius of gyration was calculated in MDAnalysis for each peptide chain/segment using Cα atoms only (name CA) with AtomGroup.radius_of_gyration().

To assess chain-level conformational variability, we then calculated the distribution of the *R*_!_ across all monomers for each structure and visualized these distributions using kernel density estimation (KDE) to compare structural variability across monomer copy numbers and sequence variants (**SI Eq. 2**). KDE provides a continuous representation of size distribution, enabling assessment of structural heterogeneity and compaction trends across different sequence variants and assembly sizes. KDE plots were generated in Python using Seaborn kdeplot. Unless otherwise specified, Seaborn’s default Gaussian KDE bandwidth rule was used and multiplied by bw_adjust=0.5.

We calculated atomic packing efficiency using PyRosetta’s packstat metric, calculated with pyrosetta.rosetta.core.scoring.packstat.compute_packing_score(pose), which is independent of scoring functions. We computed packing scores for each predicted assembly and compared across different monomer counts to assess changes in core packing and structural organization (**SI Eq. 3**). Higher values indicate tighter, void-free packing characteristic of stable folded cores, while lower values reflect under-packed or loosely organized structures. Generally speaking, proteins with less than 2,000 residues had a mean interior packing density of 0.74 and poorly packed proteins had densities less than 0.71.^77^

We quantified the assemblies’ global shape through the multimetric structures’ aspect ratios (**SI Eq. 7**) using a principal-axis-based method. Higher aspect ratios indicate elongated or fibrillar assemblies, whereas lower values indicate more compact, isotropic, or spherical morphologies. Because aspect ratio is insensitive to absolute size, it enables comparison of aggregate shape across assemblies containing different numbers of monomers. As such, it provides a complementary measure to distance- and packing-based metrics for assessing how assembly morphology evolves with increasing copy number. Together, these considerations allow for systematic, quantitative evaluation of AF3-predicted peptide assemblies across sequence variants and copy numbers.

We generated residue-residue contact maps for each predicted assembly using PyRosetta to analyze structural organization and inter-residue connectivity. For each assembly, the coordinates of α-carbon atoms were extracted from the pose and residues were considered to be in contact when the distances between their α-carbons were within 8.0 Å. Contact maps were generated as symmetric binary matrices, where a value of 1 indicates a residue pair in spatial proximity and 0 indicates the absence of a contact. These maps were calculated for each sequence design across increasing oligomeric copy numbers (10, 50, 100, 150, 200, and 250 copies) to evaluate how assembly size influences residue packing, long-range interactions, and overall structural organization. Contact maps were visualized as heatmaps, where increased contact density reflects greater structural connectivity and compactness within the predicted assemblies. Reduced contact density suggests more open or loosely packed assemblies.

We analyzed the AF3-predicted assemblies as finite structures as returned by the server. We did not impose any translational, helical, or rotational symmetry, nor did we apply periodic boundary conditions. Therefore, the radius of gyration, aspect ratio, and packstat values describe the finite predicted assemblies rather than infinite or periodically repeating structures. In addition, we calculated all quantitative metrics for the single chosen AF3 output model per sequence condition. For future studies, all five AF3 outputs of the sequence condition should be analyzed and averaged to provide a better representation of the performance.

## Results and Discussion

### X9E9 phase diagrams illustrate the relationship between hydrophobicity, peptide copy number, and assembly morphology, directly reflecting effects of the hydrophobic effect, steric constraints, and residue propensity

To evaluate how residue hydrophobicity and peptide copy number govern peptide self-assembly, we designed a panel of X9E9 sequences inspired by protein-based block copolymers^78^ and elastin-like polypeptides.^79^ We varied X across hydrophobic residues (V, I, F, L, A, P) and used AlphaFold3 to assemble increasing copy numbers of each sequence. We mapped the resulting morphologies onto a phase diagram that plots side-chain hydrophobicity (by the Kyte and Doolittle scale^80^) versus peptide copy number (not to be interpreted as a concentration scale) **(Figure 1)**. Less hydrophobic amino acids (P and A) produced disorganized, inverted structures, that is, with the weakly hydrophobic residues exposed to solvent. Increasing hydrophobicity led to micelles with well-defined hydrophobic cores, and ultimately to layered β-sheets with loops. V9E9 and I9E9 formed repeating β-sheets, consistent with their known high intermolecular H-bonding propensities,^81–83^ showing interpeptide H-bonds linking strands.^83–86^ In contrast, bulkier residues (F) contributed to incomplete, saddle-like micellar structures. Beyond 100 copies, the β-sheets of F9E9 disrupt uniform β-stacking and cause curvature, perhaps due to steric hindrance from phenylalanine’s bulky, aromatic benzyl side chain.^87^ In contrast, peptides with A and L show assembly with helical peptide conformations, reflecting their high ɑ-helical propensity.^82,88^ The glutamic acid block often mimics the secondary structure of the hydrophobic block: it forms a β-strand with the F blocks, but is helical with the A blocks. Additionally, glutamic acid readily forms helices in environments where electrostatic repulsion is reduced, such as within densely packed hydrophobic cores, evident by predicted structures for A9E9 and L9E9.

**Figure 1.**
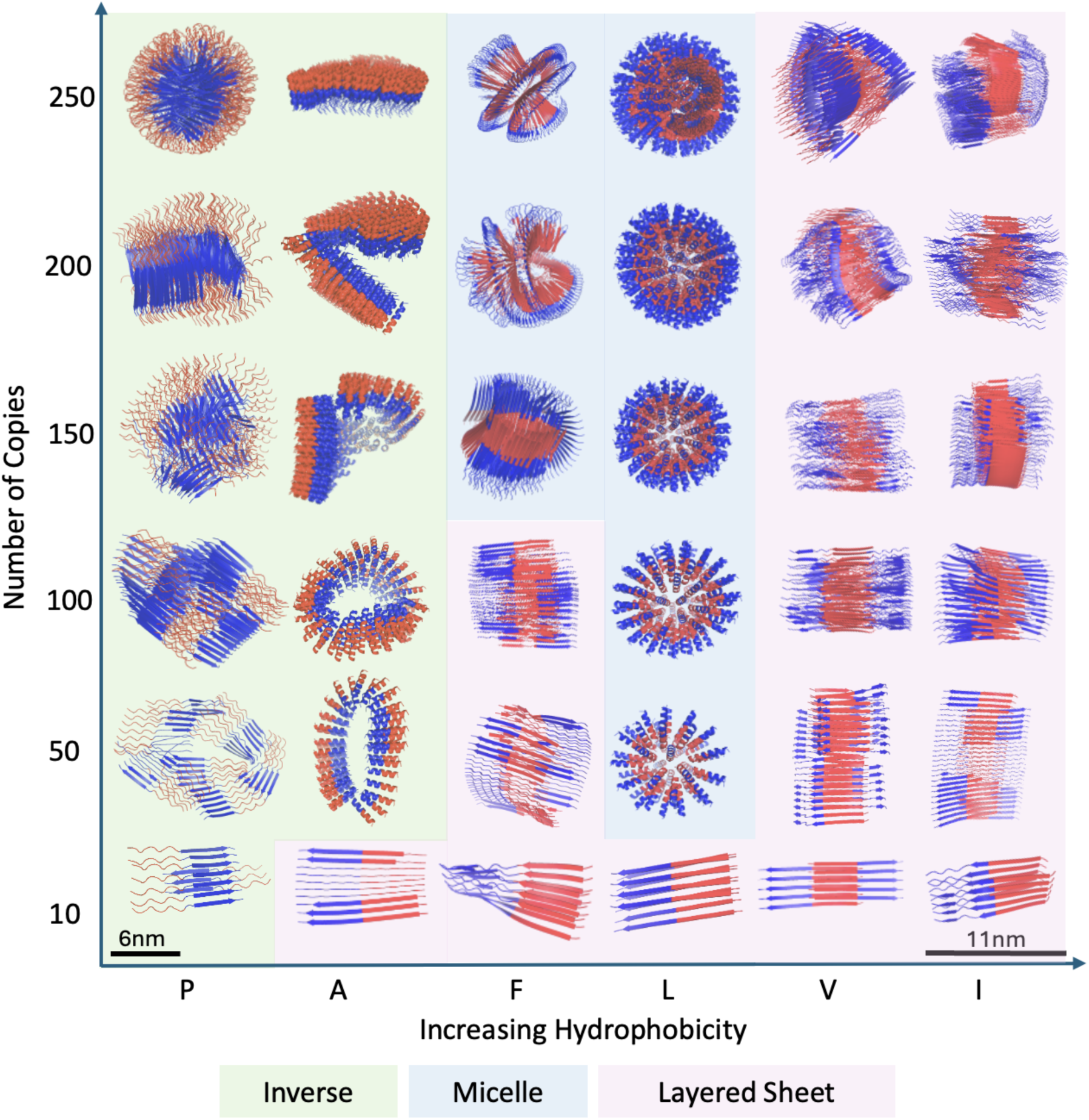
Phase diagram showing increasing hydrophobicity (by the Kyte and Doolittle scale) v.s. number of copies of X_9_E_9_ sequences, where X represents a hydrophobic amino acid. The hydrophobicity scores for the amino acids are -1.6 for proline, +1.8 for alanine, +2.8 for phenylalanine, +3.8 for leucine, +4.2 for valine, and +4.5 for isoleucine.^80^ X is colored red, while hydrophilic glutamic acid is colored blue. The diagram includes scale bars to account for the diameter of AlphaFold3’s prediction. Peptide hydrophobicity and copy number drive morphological transitions from disordered structures to micelles and layered β-sheets.

We next asked whether quantitative shape descriptors support the qualitative phase trends, and we further probe hydrophobicity effects on compactness. We examined the distribution of monomer radii of gyration (**Figure 2A**) for each X9E9 assembly (250 copies per variant) to compare how different hydrophobic residue choices influence chain compactness. L9E9 shows a narrow peak in its distribution, indicating its monomers adopt consistently compact conformations with minimal structural variability. Increasing hydrophobicity correlated with shorter radii of gyration, reflecting enhanced compactness in chain organization. These distributions highlight the uniformity within each monomer set and serve as an intuitive metric for assessing the average packing tightness, represented by the mean distance of atoms from the chain’s center.

**Figure 2.**
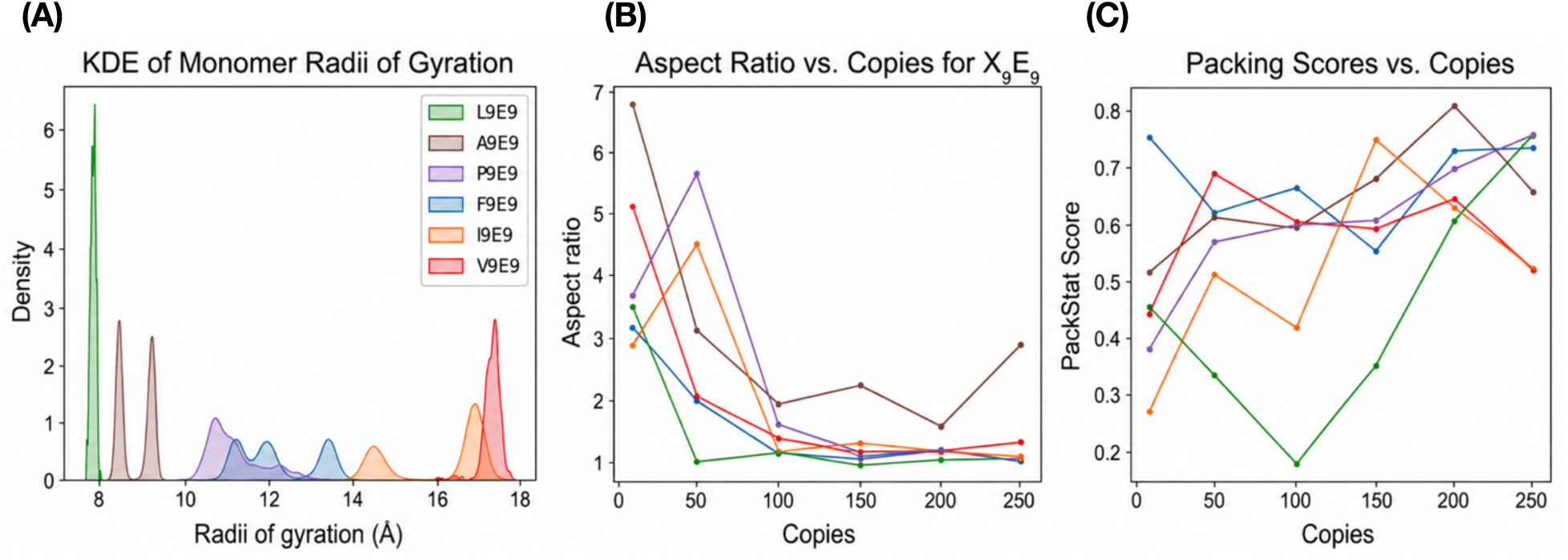
(A) **Kernel density estimation (KDE) plot of the radii of gyration** of monomers in X_9_E_9_ assemblies. Compactness increases with residue hydrophobicity; L_9_E_9_ is the most compact and morphologically uniform. (B) **Aspect ratio of structures vs number of copies** for X_9_E_9_ sequences. Aspect ratio decreased as copy number increased, indicating increased compactness. (C) Packing efficiency of X_9_E_9_ peptide sequences as a function of the number of copies of peptides modeled. Single residue changes modulate peptide packing: proline rigidity and glutamate charge drive order; side chain size controls compactness.

To quantify changes in aggregate morphology with increasing copy number, we compared aspect ratios of X9E9 assemblies (**Figure 2B**) and packing scores (**Figure 2C**) across AF3-predicted multimers. As copy number increased, aspect ratios and packing scores decreased, indicating a transition from elongated assemblies at low copy numbers toward more compact, isotropic structures.

We extended this approach to 8-block alternating sequences to evaluate whether placing residues at termini versus mid-chain changes assembly outcomes **(Figure S2)**. Qualitatively, at 10 copies, we again observed sheet-like structures, while above 50 copies, assemblies diversified into micelles, donuts, sheets, and nanotubes, with distinct outcomes depending on whether hydrophobic residues occupied termini or mid-chain positions **(Figure S2)**. Aspect ratios for these sequences showed a similar trend as that observed in X9E9 assemblies (**Figure S3**), indicating that the compaction with copy number generalizes across sequence architectures.

Together, these results suggest that AF3 often reflects residue-level propensities learned from protein structures during peptide assembly, supporting biophysical reasoning including the hydrophobic effect, steric constraints, and residue propensity. As such, these predictions should not be interpreted as a substitute for ensemble or solvent-explicit simulations. Furthermore, although the radius of gyration and aspect ratio provide quantitative measures of aggregate size and morphology, these structural descriptors alone cannot be used to predict the critical aggregation concentration (CAC). Estimation of CAC requires equilibrium measurements across known concentrations, together with an explicit system volume to relate peptide copy number to concentration. AlphaFold3 does not provide such thermodynamic information so the present analyses are limited to comparisons of aggregate morphology rather than quantitative predictions of aggregation thermodynamics.

### AF3 predictions suggest that secondary structure propensity, such as β-sheet propensity for valine and isoleucine and ɑ-helix propensity for alanine, may contribute to the predicted secondary structures of the peptide assembly

Further analyzing the X9E9 assemblies in **Figure 1**, we noted that most hydrophobic residues packed inside the structure and hydrophilic residues were solvent-facing, reflecting the hydrophobic effect of self-assembly behavior. Also, the predicted secondary structure varied with residue identity: V9E9 and I9E9 consistently formed laterally stacked β-sheets, whereas A9E9 adopted α-helical conformations. Consistent with this interpretation, residue–residue contact-map analyses reveal differences in local energetic contributions and structural connectivity among the predicted assemblies **(Figures S8-13)**. These differences may be influenced in part by the known secondary structures of the residues. In Chou and Fasman’s propensity scale,^82^ V and I have high β-sheet propensities (0.282 and 0.274, respectively), which explains their preference for β-strand conformations in AlphaFold3 predictions. In contrast, A has a substantially higher α-helix propensity (0.522) compared to its β-sheet propensity (0.167), accounting for the helical structures observed in A9E9 assemblies. However, secondary structure propensity is only one of several factors that may influence peptide assembly morphology. Other contributions, including steric effects, aromatic interactions, and terminal charge interactions, may also influence the observed structural organization. Thus, AlphaFold3 predictions reflect not only the global organization driven by global physiochemical properties but also the local secondary structure preferences encoded by individual residue properties.

To validate trends in secondary structure, we looked to the study of Gonzalez et al. (2024) that used CD and TEM to characterize peptide assemblies and secondary structure.^89^ They observed that FRV3FR self-assembled into nanofibrils and ultimately macroscopic hydrogels, possessing β-sheet secondary structure and evidence of aromatic stacking from CD characterization. They did not report any existing secondary structures in FRA3FR. In AlphaFold3 simulations, FRV3FR uniformly formed β-sheets (**Figure 3A**), in agreement with Gonzalez et al’s observations. However, FRA3FR in AlphaFold3 aggregates into helical structures (**Figure 3B**), in contrast to experimental observations.

**Figure 3.**
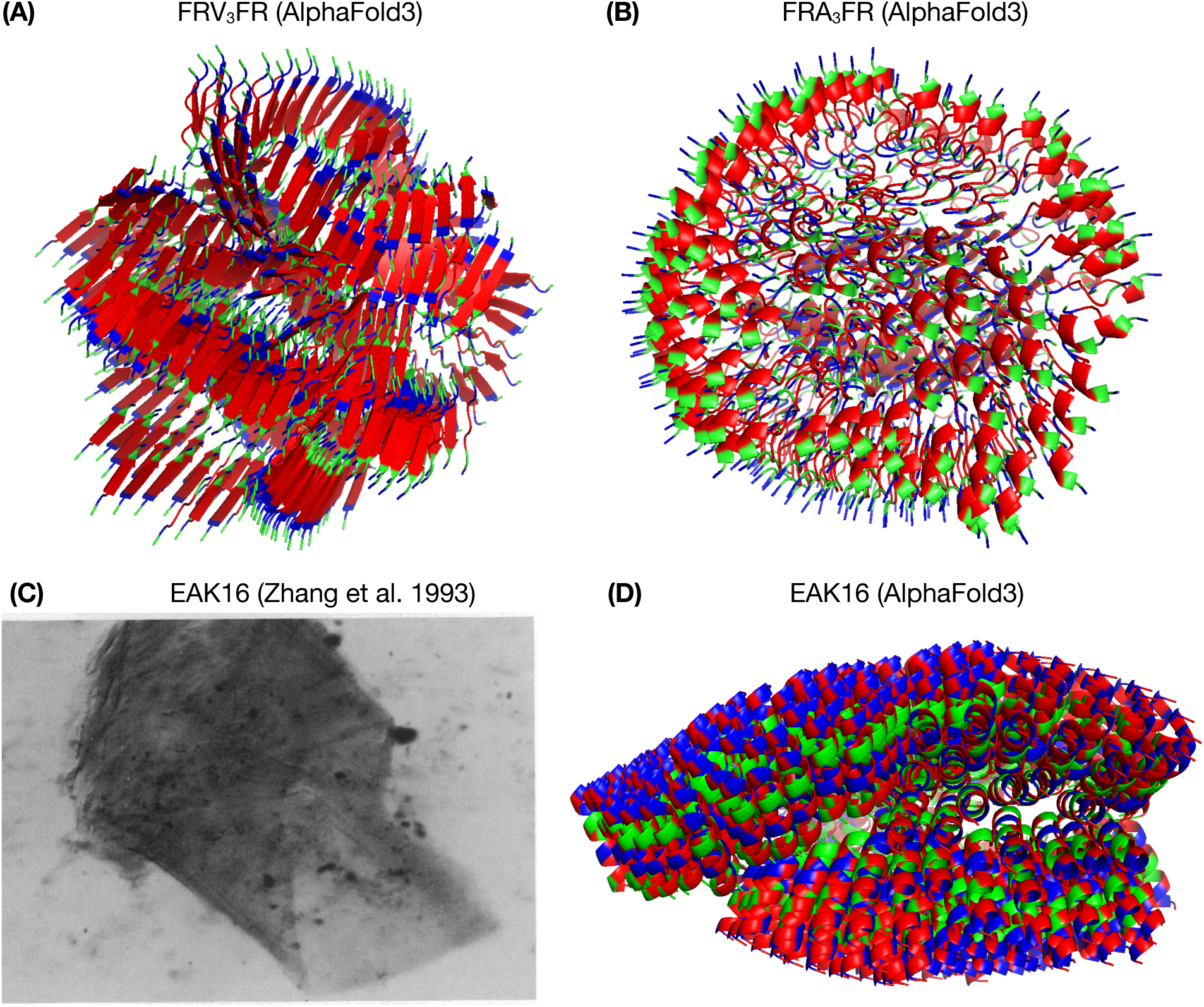
(A) **AlphaFold3 Simulations of 500 copies of FRV_3_FR**, capturing β-sheet secondary structure. Valine is depicted as red, arginine is blue, and phenylalanine is green. (B) **AlphaFold3 Simulations of 500 copies of FRA_3_FR**, capturing non-β-sheet secondary structure. Alanine is depicted as red, arginine is blue, and phenylalanine is green. (C) **Experimental TEM image for the self-assembled EAK 16 membrane** in water after the addition of salt. Reprinted from Zhang et al. Copyright 1993 National Academy of Sciences. (D) **300 copies of EAK16** arranged per AlphaFold3’s prediction, where alanine is red, lysine is green, and glutamic acid is blue.

In the case of EAK16, or (AEAEAKAK)2, experimentally studied by Zhang et al (1993),^40^ the AlphaFold3-predicted secondary structure did not match the experimental findings. Zhang et al. reported that EAK16 has a characteristic β-sheet circular dichroism spectrum and self-assembles into a membrane in water upon the addition of salt (**Figure 3C**). AlphaFold3 predicted the supramolecular membrane structure correctly, but incorrectly predicted an ɑ-helical structure (**Figure 3D**) rather than β-sheet. We speculate that AlphaFold3 prioritized alanine’s high ɑ-helical propensity over the interactions that drive β-sheet formation experimentally. Amphiphilic peptides often reorganize into helices when packing requires curvature, prioritizing helical conformations in densely packed assemblies.^77,88^

### An expanded design of alternating V and E sequences shows that longer hydrophobic segments lead to disordered structures, while longer hydrophilic segments promote compactness and organization

To investigate how segment length impacts assembly, we varied the number of valines (hydrophobic sequence length) in amphiphilic molecular designs: V2E2, V4E2, V6E2, V8E2. As segments lengthened, predicted assemblies showed reduced global order (**Figure S4**). Longer sequences (e.g., V6E2 and V8E2) exhibited surface exposure of hydrophobic residues without clear hydrophilic-hydrophobic segregation, lacking the compartmentalization typical of amphiphilic assemblies. The exposure of hydrophobic residues likely indicates physical packing challenges rather than a realistic stable structure, highlighting a limitation of AF3 in extrapolating to systems where the sequence balance strongly disfavors ordered self-assembly. At the chosen high copy number, AF3 fails to generate a physically meaningful structure, possibly because of its predisposition to predicting structures as one assembly, rather than multiple units. As copy number drives the prediction of the correct self-assembled structure, iterating copy number for a specific system may prove to be a useful strategy for prediction or design.

We next varied hydrophilic segment length (V2E2, V2E4, V2E6, and V2E8) while keeping the valine numbers constant. Increasing hydrophilic residues led to progressively more compact and ordered assemblies (**Figure S5**). V2E4 showed partial hydrophobic exposure, while V2E8 formed a well-organized structure with hydrophobic cores buried and hydrophilic regions solvent exposed. Longer hydrophilic segments tend to enhance solvation and drive more complete hydrophobic packing, suggesting that AF3’s predictions reflect physically expected behavior.

In comparison to the other sequences in **Figure 3** and **Figure 4**, the tetrapeptide V2E2 is particularly poorly defined. This difficulty with tetrapeptides likely arises from limitations in AF3’s training and its focus on static structures rather than dynamic ensembles. Four amino acids is AF3’s minimum input for a peptide length, suggesting that is the lower limit of successful predictive ability. Given that AlphaFold models are predominantly trained on longer, folded globular proteins, they struggle with the shorter sequences in our study, resulting in lower predictive confidence. Minimalist peptides are inherently challenging to design experimentally because their noncovalent interactions are relatively weak, requiring high concentrations (and thus close molecular proximity) to overcome kinetic barriers and achieve stable, organized assembly.^45^ We suggest that AF3’s maximum copy number for tetrapeptides may be insufficient to capture this threshold behavior and accurately predict assembly formation.

**Figure 4.**
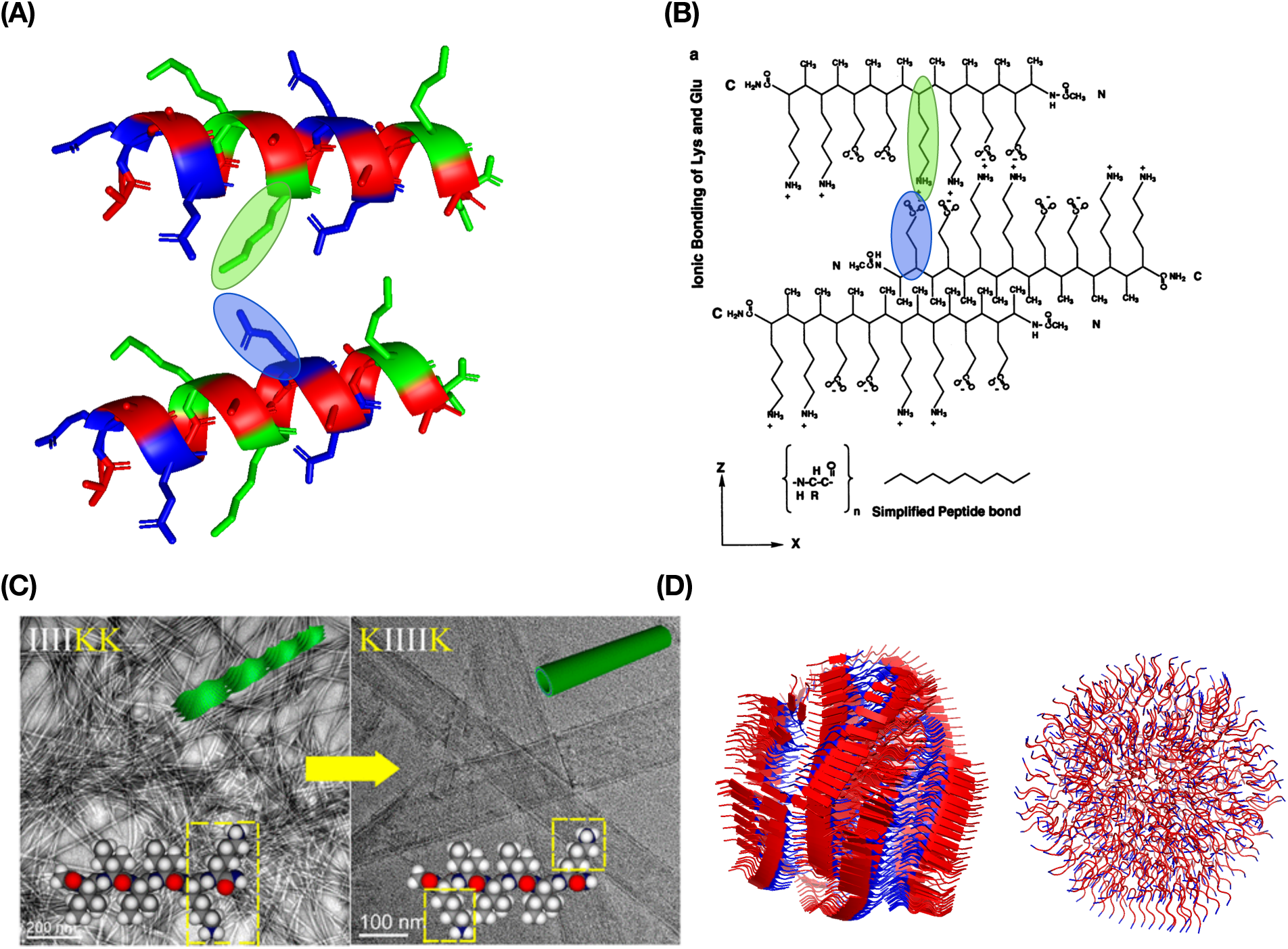
(A) **Two individual peptides selected from AF3’s structural prediction of EAK16**, emphasizing that charged lysine and glutamic acid side chains are positioned facing each other to form ionic bonds. Alanine is red, lysine is green, and glutamic acid is blue. (B) **Schematic depicting ionic bond formation and side chain arrangement of individual EAK16 peptides**. Reprinted from Zhang et al. Copyright 1993 National Academy of Sciences. (C) **TEM images of nanoscale structures**. IIIIKK shows thin, twisted filaments while KIIIIK shows wide, ribbon-like nanobelts due to less electrostatic repulsion and greater opportunity for lateral stacking. Reprinted (adapted) with permission from Zhao et al., *Langmuir* **29** (44), Copyright 2013 American Chemical Society. (D) **AF3 depiction of the same assemblies**, modeling 500 copies of each peptide, IIIIKK on the left and KIIIIK on the right. Isoleucine is red, lysine is blue. AF3 accurately predicts the secondary structure for IIIIKK but struggles with predicting the correct structure when the lysine residues are positioned close to isoleucine residues.

Additionally, the absence of explicit solvent modeling and thermodynamic sampling may further limit AF3’s ability to predict the true packing patterns of amphiphilic systems. The model does not incorporate concentration or environmental parameters. Instead, users specify only the number of monomer copies included in a prediction. While increasing the number of copies can be used to explore potential assembly geometries, the specified copy number should not be interpreted as a physical concentration or directly related to the critical aggregation concentration (CAC). Unlike traditional molecular dynamics or Monte Carlo simulation methods, which can show a multi-component system with separate complexes in one simulation box.^71^ AF3 predicts a single assembled structure containing all provided monomers. It does not consider whether they might naturally separate into smaller assemblies based on steric limitations or concentration-dependent behavior, potentially leading to strained or unrealistic packing arrangements. As a result, the predicted structures may not fully capture the concentration-dependent equilibria and dynamic self-assembly behavior characteristic of amphiphilic peptide systems.

To further test the effect of alternating hydrophobic and hydrophilic patterns, we designed several longer sequences. We tested the effect of larger blocks of V and E, designing a two-block sequence (e.g. V9E9) and an 8-block alternating sequence (e.g. (V4E4)4 ) (**Figure S6**). Compared to tetrapeptides, both designs showed improved self-assembly, with hydrophobic cores and hydrophilic shells. Repeating blocks promoted formation of nano-doughnut structures, contrasting with the sheet-like aggregates of the two-block sequence and the disordered micelles seen with tetrapeptides. In the supplement, we also assessed how sequence inversion shifted the spatial distribution of key intermolecular interactions, disrupting cooperative packing and impacting the formation of a thermodynamically stable core (**Figure S7**).

These results underscore the importance of segment length and block architecture in dictating peptide assembly morphology. AF3 reliably captures the trend toward improved order and stability with longer hydrophilic segments and alternating block patterns, producing well-defined hydrophobic cores and hydrophilic shells. However, its limitations become apparent at short peptide lengths (4 amino acids) and for sequences with extreme hydrophobicity (80% hydrophobic residues), reflecting the constraints of its training on static structures in deep energy wells and the lack of modeling dynamics and solvent. Future studies must use MD strategies or BioEmu to generate ensembles for dynamical protein design with an understanding of equilibrium distributions of both protein folding and native-state conformational transitions.^91^

### AF3 captures electrostatic preferences when predicting self-assembly, but struggles with long distance electrostatic patterning when charged residues are positioned at opposite termini

To probe the effects of charged residues, we probed deeper into the EAK16 case from Zhang et al (1993).40 They described that the β-sheet is stabilized by hydrophobic interactions between opposing A side chains on one face, while K and E side chains on the opposite face form ionic bonds (**Figure 4B**). To examine the microstructure predicted by AF3, we extracted two chains from the supramolecular complex (**Figure 4A**). Consistent with Zhang et al., we observe alanine side chains clustered together and K-E salt bridges; however, AF3 predicts a helical conformation rather than the experimentally observed β-sheet structure. Thus, although AlphaFold3 can generate local arrangements of residues based on opposing charges, it does not imply that it quantitatively models the energetics of the assembly.

More generally, AlphaFold3 only implicitly captures charge-based structural preferences. The model may learn sequence-structure correlations involving charged residues from its training data, but it does not explicitly incorporate effects from solvent, ionic strength, or pH. Therefore, the predicted electrostatic patterns should be treated as structural hypotheses rather than explicit predictions of electrostatic or pH/salt-dependent assembly behavior.

We additionally compared AlphaFold3 predictions to the study by Zhao et al. (2013) describing different 1D nanostructures resulting from small sequence rearrangements in peptides composed of K and I.47 Their TEM data showed that IIIIKK formed thin nanofibrils and KIIIIK formed wide nanotubes (**Figure 4C**). For IIIIKK, AF3 captures the presence of β-sheet secondary structure and predicts a filament-like morphology, consistent with the experimentally observed thin fibrils (**Figure 4D**). However, for KIIIIK, AF3 fails to reproduce the correct secondary structure or higher-order morphology, instead generating assemblies that lack the extended, laterally stacked organization characteristic of the experimentally observed nanotubes.

Zhao et al. attributed these divergent assembly behaviors to differences in the balance between hydrophobic interactions among isoleucine residues and electrostatic repulsion between charged lysine side chains. In IIIIKK, adjacent lysine residues at the C-terminus experience greater electrostatic repulsion, limiting lateral stacking and promoting twisting. These twisted ribbons evolve into helical ribbons and eventually nanotubes, which is experimentally consistent with principles of hierarchical self-assembly.^92^ On the other hand, the spatial separation of lysine residues reduces electrostatic frustration at the sheet-sheet interface, allowing for the formation of wider, ribbon-like nanostructures. AF3’s failure to capture this behavior may arise from difficulty in resolving long-range electrostatic patterning, particularly when charged residues are positioned at opposing termini. Because AlphaFold3 does not explicitly compute energies driving electrostatic repulsion, ionic screening, or solvent interactions, this misprediction should be interpreted as a limitation in the model’s learned structural representation, rather than a failure of an energy model.

We further investigate how AF3 captures electrostatic forces by examining peptides developed by Dong et al. (2007), where the self-assembly was attributed to molecular frustration.41 Monomers were designed to follow an ABA motif. The A-block is one or multiple charged residues (e.g., K) to improve solubility and interact with salt in solution, and the B-block consists of varying numbers of units of alternating hydrophobic (L) and hydrophilic (Q) residues, specifically alternating so as to produce distinct hydrophilic and hydrophobic molecular faces (**Figure 5A**). AF3 confirms this monomer conformation in ABA assemblies (**Figure 5B**).

**Figure 5.**
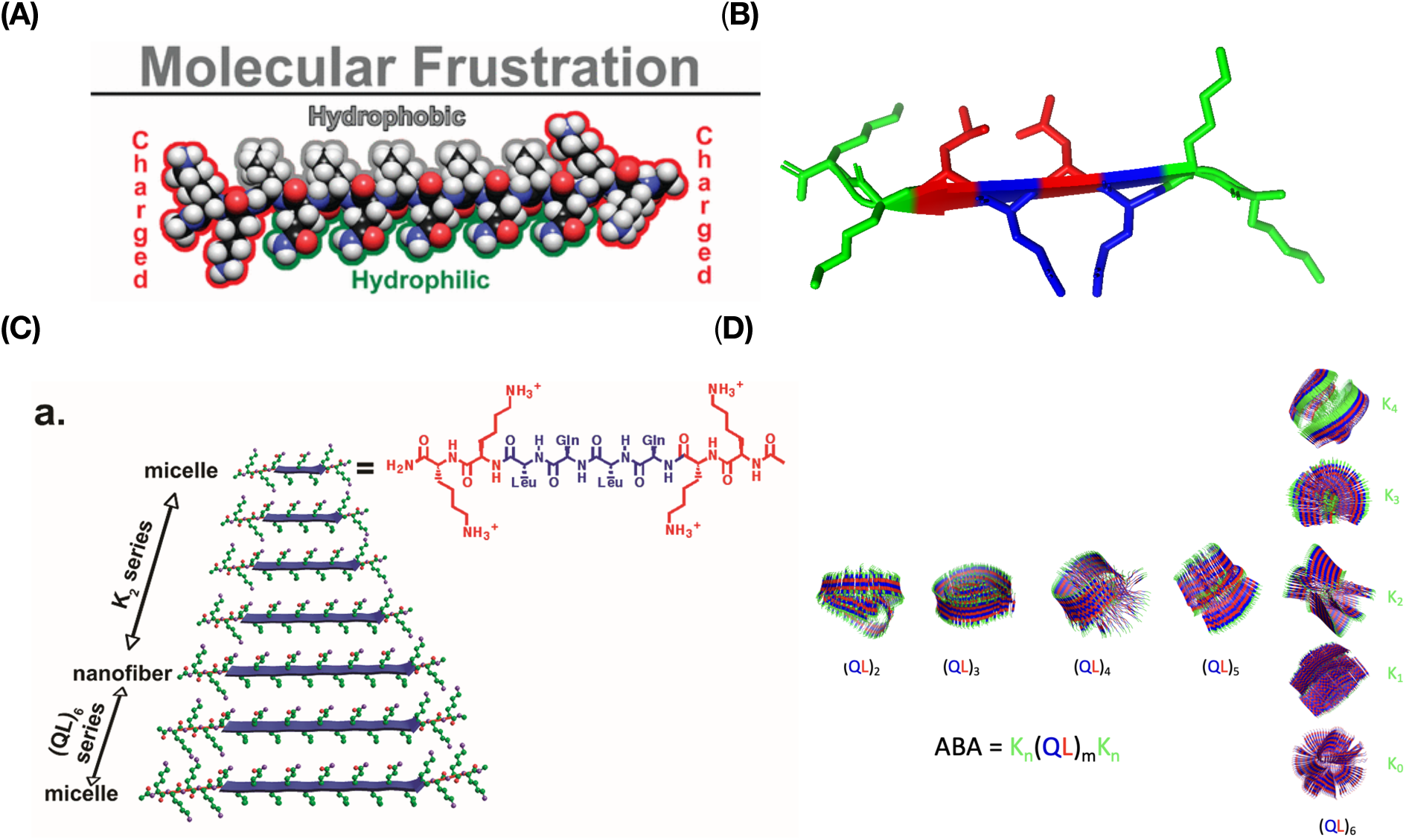
(A) **Schematic illustrating molecular frustration**. Reprinted (adapted) with permission from Dong et al., *JACS* **129** (41). Copyright 2007 American Chemical Society. Hydrophobic and hydrophilic residues are alternating (varying repeating units) such that they are positioned on opposite sides of the peptide, which is capped with charged residues (varying number). (B) **AF3 depiction of a single monomer in an assembly (n=2, m=2)**, validating the existence of the hydrophobic and hydrophilic faces on opposite sides of the molecule. Lysine is green, glutamic acid is blue, and leucine is red. (C) **Schematic illustrating experimental design and self-assembly trends with molecular modifications**. Reprinted (adapted) with permission from Dong et al., *JACS* **129** (41). Copyright 2007 American Chemical Society. As the number of repeating units of QL (m) increases and the number of lysines (n) is held constant at 2, the structures shift from micellar to nanofibers. As the number of lysines increases and the repeating units of QL are held constant to 6, the structures shift back to micelles. (D) **AF3 depiction of the same schematic**, running 250 copies of each experimentally determined sequence.

Dong et al. systematically varied the number of K’s in the A-block and the number of (QL)’s in the B-block. As the B-block increased in length (repeating units), the structure shifted from micellar to nanofibrillar. As the charged A-block increased in length, the structure shifted back to micellar (**Figure 5C**). Replicating this experiment in AF3, we observed similar trends (**Figure 5D**). As the number of alternating QL dimers (length of B-block) increases, the structure becomes more fibrillar. As the number of K residues in the A-block gets farther from 2, it becomes more micellar. This trend is observable in either direction (from 2 to 0, or 2 to 4), but as the number of K residues increases there is a change in side chain orientation where the K residues tuck slightly inwards, facing each other, which may contradict their experimental role in increasing solubility, charge screening, and overall solvent interactions. Again, this inconsistency shows that AlphaFold3 can identify structural patterns associated with charged sequence blocks, but it does not explicitly model the solution environment.

To further examine AF3’s interpretation of charged amino acids at the termini of sequences, we investigated a series of IKVAV-containing peptides originally designed by Rodriguez et al. (2013) with increasing numbers of of N-terminal D residues: IKVAV, DIKVAV, DDIKVAV and DDDIKVAV.^93^ They found incorporation of additional D residues systematically lowered the overall pKa of the peptides, thereby modulating the pH responsiveness of the system. All sequences were found by TEM to self-assemble into nanofibers with consistent β-sheet secondary structure, yielding nanostructured fibrous networks with broadly comparable morphologies. Notably, the network formed by DDDIKVAV was more fragmented than those of the shorter sequences, suggesting that four aspartic acid residues represent the upper limit of structural stability for this system.

We replicated this series in AF3 (**Figure 6**). At a copy number of 500, AF3 is unable to accurately capture the filamentous morphology or the consistent β-sheet secondary structure exhibited by these peptides, instead showing a liquid condensate-like morphology devoid of any secondary structure (**Figure 6**, right column). To determine whether this discrepancy arose from AF3’s interpretation of residue-specific properties and charge states, or from another limitation, we performed additional simulations at 50, 150, and 250 copies (**Figure 6**). Predicted assemblies improved (correct β-sheet prediction) as copy number decreased, aligning with the CD-validated β-sheet structure. This finding underscores that maximizing copy number is not inherently beneficial in AF3; instead, accurate predictions require identifying a system-specific optimum through iterative tuning.

**Figure 6.**
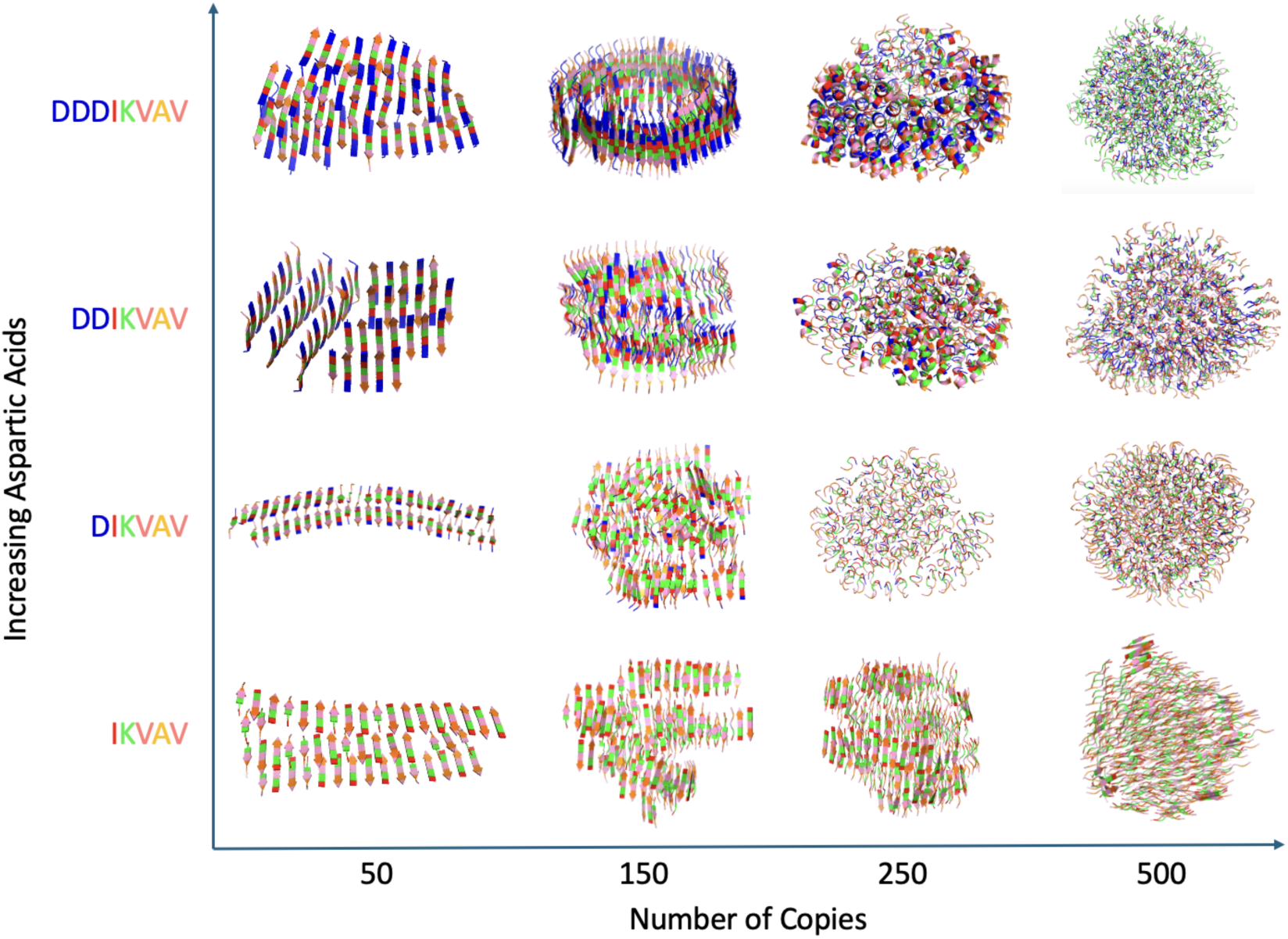
AF3 simulation of terminally charged peptides designed by Rodriguez et al. (2013). 50, 150, 250, and 500 copies of each peptide: IKVAV, DIKVAV, DDIKVAV, DDDIKVAV. Isoleucine is red, valine is pink, alanine is orange, lysine is green, and aspartic acid is blue. AF3’s ability to predict the correct CD-validated β-sheet structure improves with decreasing copy number.

A limitation for short, charged peptides is that they are subject to charge regulation based on the pH, neighboring proteins and ligands, and other factors affecting its protonation state. This phenomenon influences self-assembly and hydrogel behavior by affecting the proton equilibria of charged sites.^94,95^ Since AlphaFold3 does not explicitly represent such charge-fluctuating effects, its predictions for peptides with clusters of acidic/basic residues should be interpreted as a hypothesis rather than a prediction of pH-dependent assembly behavior.

To summarize, Table 1 provides a comprehensive, experiment-by-experiment comparison of AlphaFold3 predictions with experimentally characterized peptide self-assembly systems, with each entry evaluated based on predicted secondary structure, supramolecular morphology, hydrophobic assembly, and electrostatic interactions.

**Table 1.** Comparison of AlphaFold3 predictions with experimentally reported peptide self-assembly behavior. Agreement (✓, ✕) was assessed based on predicted secondary structure, supramolecular morphology, hydrophobic assembly, and electrostatic interactions relative to the corresponding experimental studies. “Partial” indicates that AlphaFold3 correctly captured one or more aspects of the assembly process but failed to reproduce all experimentally observed structural features.

| Peptide Sequence<br>(Reference) | Copy<br>No. | Secondary<br>Structure | Morphology /<br>Aggregation | Electrostatics | Overall<br>Agreement |
| --- | --- | --- | --- | --- | --- |
| FRV<sub>3</sub>FR (Gonzalez et al.<sup>89</sup>) | 500 | ✓ β-sheet | ✓ Fiber | – | Yes |
| FRA<sub>3</sub>FR (Gonzalez et al.<sup>89</sup>) | 500 | ✗ α-helix | ✗ Aggregated | – | No |
| EAK16 (Zhang et al.<sup>40</sup>) | 300 | ✗ α-helix | ✓ Membrane<br>assembly | ✓ Ionic interactions | Partial |
| IIIIKK (Zhao et al.<sup>47</sup>) | 500 | ✓ β-sheet | ✓ Fiber | ✓ Repulsion due to<br>proximal lysines | Yes |
| KIIIIK (Zhao et al.<sup>47</sup>) | 500 | Partial | ✗ Limited lateral<br>growth | ✗ Long range patterning | No |
| K<sub>2</sub>(QL)<sub>2</sub>K<sub>2</sub> (Dong et al.<sup>41</sup>) | 250 | ✗ β-sheet | Partial | – | Partial |
| K<sub>2</sub>(QL)<sub>3</sub>K<sub>2</sub> (Dong et al.<sup>41</sup>) | 250 | ✗ β-sheet | Partial | – | Partial |
| K<sub>2</sub>(QL)<sub>4</sub>K<sub>2</sub> (Dong et al.<sup>41</sup>) | 250 | ✗ β-sheet | Partial | – | Partial |
| K<sub>2</sub>(QL)<sub>5</sub>K<sub>2</sub> (Dong et al.<sup>41</sup>) | 250 | ✓ β-sheet | ✓ Fiber | – | Yes |
| K<sub>2</sub>(QL)<sub>6</sub>K<sub>2</sub> (Dong et al.<sup>41</sup>) | 250 | ✓ β-sheet | ✓ Fiber | – | Yes |
| (QL)<sub>6</sub> (Dong et al.<sup>41</sup>) | 250 | ✓ β-sheet | ✗ Micelle | Partial | Partial |
| K(QL)<sub>6</sub>K (Dong et al.<sup>41</sup>) | 250 | ✓ β-sheet | ✓ Fiber | Partial | Partial |
| K<sub>3</sub>(QL)<sub>6</sub>K<sub>3</sub> (Dong et al.<sup>41</sup>) | 250 | ✓ β-sheet | ✓ Micellar<br>transition | ✓ Charge effects of<br>added lysines | Yes |
| K<sub>4</sub>(QL)<sub>6</sub>K<sub>4</sub> (Dong et al.<sup>41</sup>) | 250 | ✗ β-sheet | Partial | Partial charge effects of<br>added lysines | Partial |
| IKVAV (Rodriguez et al.<sup>93</sup>) | 50 | ✓ β-sheet | ✓ Fiber | – | Yes |
| IKVAV (Rodriguez et al.<sup>93</sup>) | 150 | ✓ β-sheet | ✓ Fiber | – | Yes |
| IKVAV (Rodriguez et al.<sup>93</sup>) | 250 | ✓ β-sheet | ✓ Fiber | – | Yes |
| IKVAV (Rodriguez et al.<sup>93</sup>) | 500 | ✓ β-sheet | ✓ Fiber | – | Yes |
| DIKVAV (Rodriguez et al.<sup>93</sup>) | 50 | ✓ β-sheet | ✓ Fiber | – | Yes |
| DIKVAV (Rodriguez et al.<sup>93</sup>) | 150 | ✓ β-sheet | ✓ Fiber | – | Yes |
| DIKVAV (Rodriguez et al.<sup>93</sup>) | 250 | ✗ Coil | ✗ Micelle | – | No |
| DIKVAV (Rodriguez et al.<sup>93</sup>) | 500 | ✗ Coil | ✗ Micelle | – | No |
| DDIKVAV (Rodriguez et al.<sup>93</sup>) | 50 | ✓ β-sheet | ✓ Fiber | – | Yes |
| DDIKVAV (Rodriguez et al.<sup>93</sup>) | 150 | ✓ β-sheet | ✓ Fiber | – | Yes |
| DDIKVAV (Rodriguez et al.<sup>93</sup>) | 250 | ✗ Helix/Coil | ✗ Micelle | – | No |
| DDIKVAV (Rodriguez et al.<sup>93</sup>) | 500 | ✗ Helix/Coil | ✗ Micelle | – | No |
| DDDIKVAV (Rodriguez et al.<sup>93</sup>) | 50 | ✓ β-sheet | ✓ Fiber | – | Yes |
| DDDIKVAV (Rodriguez et al.<sup>93</sup>) | 150 | ✓ β-sheet | ✓ Fiber | – | Yes |
| DDDIKVAV (Rodriguez et al.<sup>93</sup>) | 250 | ✗ Helix/Coil | ✗ Micelle | – | No |
| DDDIKVAV (Rodriguez et al.<sup>93</sup>) | 500 | ✗ Helix/Coil | ✗ Micelle | – | No |

**Table 2** synthesizes these observations across broader categories to identify structural and physicochemical features that AlphaFold3 predicts reliably and those that remain challenging. AlphaFold3 is most reliable when predicting local structural features governed by intrinsic peptide properties, including β-sheet formation, residue-level packing, and hydrophobically driven assembly. Prediction accuracy decreases for systems in which supramolecular organization is dominated by environmental context, long-range electrostatic patterning, solvent-dependent interactions, or concentration-dependent structural transitions. These observations suggest that AF3 is best suited for identifying intrinsic sequence-driven assembly tendencies rather than fully predicting experimental self-assembly outcomes. These sequence-driven predictions provide useful design criteria for prioritizing candidate sequences for subsequent physics-based and energetic simulations, enabling AF3 to serve as an initial screening step before more computationally intensive molecular dynamics approaches.

**Table 2:**
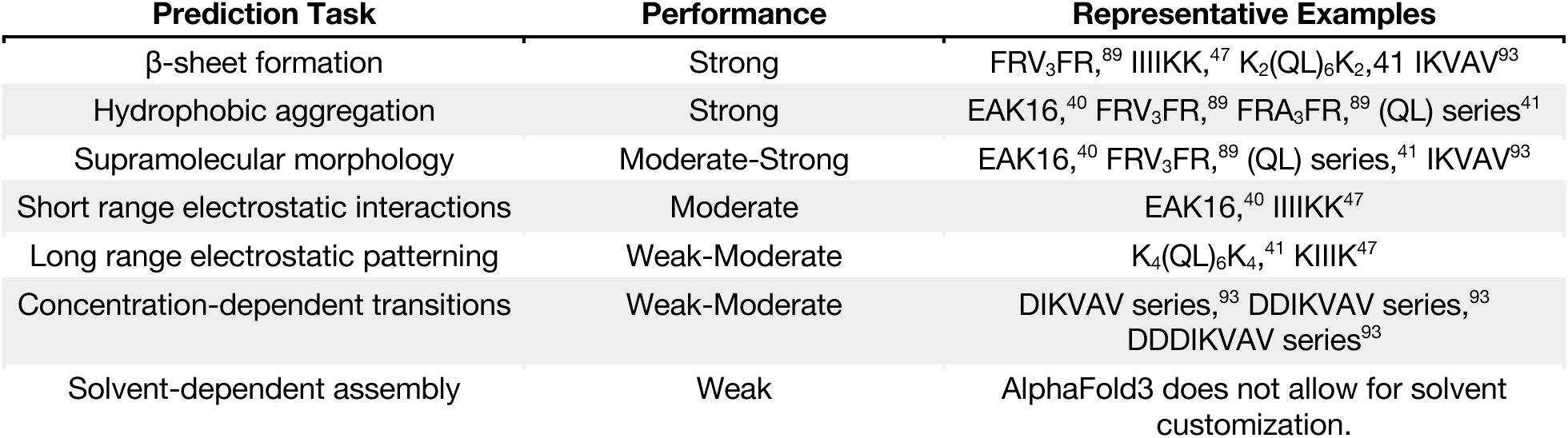
Summary of AlphaFold3’s capability of predicting specific classes of structural and physicochemical features, presented alongside representative examples.

## Conclusion

Choosing residues based on their physicochemical properties (acidity, charge, steric effects, and secondary structure propensities) strongly influences peptide self-assembly both internally and globally, as reflected in both our AF3 observations and experimental studies in the literature. Across multiple experimentally validated systems, AF3 captures key sequence-driven features of peptide self-assembly, particularly secondary structure formation, residue-level packing interactions, hydrophobic and hydrophilic patterning, and directional trends in supramolecular assembly behavior resulting from small sequence modifications. These predictions are consistent with established biophysical principles governing peptide assembly, including hydrophobic interactions, steric constraints, secondary structure propensities, and electrostatic interactions, suggesting that AF3 can provide useful insight into intrinsic sequence-dependent assembly tendencies.

Despite these strengths, AF3 remains fundamentally limited by its inability to explicitly model the environmental and dynamic factors that govern peptide self-assembly. Solvent composition, pH, ionic strength, concentration, and dynamic exchange between assembled and unassembled peptides all strongly influence experimental assembly behavior but are absent from AF3 predictions, limiting translation to systems such as the solvent-dependent assemblies reported by Dong et al.^41^ or the pH-responsive morphologies described by Rodriguez et al.^93^ Furthermore, AF3 was developed to predict predominantly static protein structures, making its application to large, dynamic peptide assemblies and very short synthetic peptides inherently challenging. Consequently, while AF3 reliably predicts many intrinsic sequence-driven assembly tendencies, its accuracy decreases when supramolecular organization is dominated by environmental context, long-range electrostatic patterning, concentration-dependent structural transitions, or system size. Importantly, our results further demonstrate that copy number should be treated as a tunable, system-specific parameter, as the predicted assemblies were often sensitive to the number of peptide chains included in the calculation. Rather than adopting a universal copy number, iterative optimization may be necessary to identify assemblies that best represent the expected supramolecular behavior.

In addition, this study is restricted to peptides consisting of the twenty canonical amino acids. This is an important limitation as many synthetic self-assembling peptides have non-natural amino acids, cyclization, or other functional groups that affect assembly behavior.^96,97^ Although AF3 may be able to represent some noncanonical chemistry through learned atom-level interactions, its accuracy for such peptide systems remains insufficiently validated. A recent study benchmarking AF3 for noncanonical cyclic peptides suggests promise but emphasizes the need for specialized datasets and experimental validation.^98^ Thus, AF3 predictions for NCAA-containing self-assembling peptides currently represent an important direction for future benchmarking.

These limitations highlight the need for predictive models developed specifically for self-assembling peptide materials rather than folded proteins. Unlike proteins that adopt stable native structures,^99^ peptide assemblies exist as dynamic supramolecular ensembles whose structures are highly sensitive to environmental conditions and continuously exchange monomers through grand-canonical assembly processes. Future models should therefore extend beyond static structure prediction by incorporating experimentally relevant contextual parameters (including solvent composition, pH, ionic strength, concentration, temperature, and copy number), together with experimental self-assembly data such as CD, FTIR, TEM, or cryo-EM. Recent computational efforts demonstrate early progress toward this goal, with physics-based simulations incorporating environmental context into peptide self-assembly prediction^100,101^ and machine-learning approaches demonstrating the feasibility of curating more than 1000 sequence-condition-assembly relationships from the literature to predict assembly phases with >80% accuracy.^102^ Building on these efforts will require larger, well-curated datasets that systematically connect peptide sequence, environmental conditions, and experimentally measured assembly outcomes. Incorporating these variables will require large, well-curated datasets that systematically connect peptide sequence, environmental conditions, and experimentally measured assembly outcomes. Unlike protein structures, for which extensive databases of experimentally determined structures exist, self-assembling peptide datasets remain limited and heterogeneous, often lacking standardized reporting of assembly conditions and quantitative structural descriptors. Systematic curation of existing and future experimental studies demonstrating the effects of peptide sequence,^31,39–54^ concentration,^39,55–57^ pH,^17,39,58–60^ and solvent composition^55,56,61–63^ on supramolecular morphology could provide a foundation for training machine-learning models to predict environment-dependent peptide self-assembly and develop transferable sequence-environment-structure relationships. Additionally, incorporating dynamic ensemble sampling approaches such as BioEmu^91^ may further improve predictive accuracy and generalizability.

While the development of new predictive models remains an important goal, existing tools such as AF3 already offer considerable value. We envision AF3 as an accessible first-pass screening tool within a hierarchical computational design workflow including molecular simulation and experimental characterization. AF3 enables rapid screening of candidate sequences, visualization of secondary structure and intermolecular packing, and hypothesis generation within minutes using pattern matching that reflects intrinsic sequence-driven features such as β-sheet propensity, hydrophobic patterning, and residue-level interactions. This speed enables evaluation of a broad range of peptide sequences and copy-number conditions, as demonstrated in this study, that would otherwise be prohibitively time-consuming to examine using simulations alone. Promising candidates can then be prioritized for more computationally intensive coarse-grained molecular dynamics simulations to investigate environmentally dependent assembly pathways, copy-number effects, and concentration-dependent behavior, followed by all-atom simulations to characterize hydrogen bonding, electrostatic interactions, and interfacial packing before experimental validation with CD and/or TEM. Accordingly, AF3 and molecular dynamics should be viewed as complementary rather than competing computational approaches, with AF3 providing rapid sequence-driven structural hypotheses and molecular simulations resolving the environmentally dependent and dynamic aspects of peptide self-assembly.

By positioning AF3 as a rapid hypothesis-generation and sequence-screening tool within a broader computational and experimental workflow, its strengths can be leveraged while avoiding overinterpretation of its limitations. Integrating AF3 with coarse-grained and all-atom molecular simulations, together with experimental validation, provides a practical framework for accelerating the rational design of self-assembling peptide materials while improving the reliability and transferability of computational predictions.

## Supporting information

Supplemental Equations and Figures

## Data and Materials Availability

All data needed to evaluate the conclusions in this paper are present in the paper and/or the Supplementary Information.

## Supporting Information

This PDF file includes:

- Equations 1-7
- Figures S1-S13

## Funding Information

The authors acknowledge the support from the following funding: National Institutes of Health R35GM141881.

## Author Contributions

S.C., J.H., and C.S. contributed equally to conceptualization, methodology, software, investigation, data curation, formal analysis, visualization, validation, and writing of the original draft, as well as review and editing of the manuscript. J.J.G. contributed to conceptualization, supervision, and review and editing of the manuscript.

## Conflicts of Interest

Authors declare no conflicts of interest.

## Notes

### Competing Interest Statement

The authors have declared no competing interest.

### Summary of Updates

We have made several substantive revisions to place our work more clearly in the context of existing computational approaches for supramolecular assembly and to clarify how AlphaFold3 can be integrated into a broader computational workflow. We have also expanded our discussion of AlphaFold3's limitations, particularly its inability to explicitly account for environmental factors such as concentration and solvent conditions.

