## Supplemental Equations and Figures for "Predicting Supramolecular Self-Assembly of Peptide Structures with AlphaFold3"

**Affiliations**:

Johns Hopkins University

3400 N. Charles Street

Baltimore MD 21218

### Supplemental Information

#### Supplemental Methods and Figures

**Radius of Gyration**

For a given chain or multimeric assembly, the radius of gyration is defined as the root-mean-square distance of all atoms from the structure’s center of mass:

$R_{g}=\sqrt{\frac{1}{N}\sum_{i=1}^{N} ||r_{i}-r_{cm}||^{2}}$	[Eq. 1]

where *N* is the number of atoms, *r*_i_ is the position of atom *i*, and *r*_cm_ is the system's geometric center-of-mass coordinate.

**Kernel Density Estimation**

Given a set of $R_{g}$’s for each assembled structure, the KDE estimates the underlying probability density function as:

$f(d)=\frac{1}{nh}\sum_{i=1}^{n} K\left( \frac{d-d_{i}}{h} \right)$	[Eq. 2]

where *K* is a smoothing kernel and *h* is the bandwidth.

**Packing Score**

For each atom, a spherical region of fixed radius is considered, and the fraction of that volume occupied by neighboring atoms is computed. These local occupancy values are then averaged over all atoms to yield a global packing score bounded between 0 and 1. The packing score was determined using the following equation:

$P_{\mathsf{score}} = 1-\frac{V_{\mathsf{void}}}{V_{\mathsf{interior}}}$	[Eq. 3]

where $P_{\mathsf{score}}$ is the packing score, $V_{\mathsf{void}}$ is the void volume and $V_{\mathsf{interior}}$ is the total interior volume.

**Deriving Aspect Ratio**

The aspect ratio was calculated based on the principal axes of the atomic coordinate distribution. All atomic Cartesian coordinates (x,y,z) were extracted from the protein structure using PyRosetta, including every atom from each residue in the pose. These coordinates were assembled into a single N×3 matrix where N is the total number of atoms. The spatial covariance matrix of the atomic coordinates was then calculated to capture the variance of atomic positions along each dimension (Eq. 4).

$C = \frac{1}{N-1}{(X-\underline{X})}^{T}(X-\underline{X})$	[Eq. 4]

where $X$ is the matrix representation of the Cartesian coordinates of the protein atoms, $\underline{X}$ is the mean coordinate vector of all atoms. This covariance matrix captures the variance of atomic positions along each spatial dimension.

Eigenvalue decomposition of *C* was performed to obtain

$Cv_{k}=\lambda_{k}v_{k}$	[Eq. 5]

where $v_{k}$ and $\lambda_{k}$ are the eigenvectors and eigenvalues corresponding to the principal axes of the protein. The eigenvalues represent the variance of atomic positions along each principal axis. The effective length, $L_{k}$, along each principal axis was defined as the square root of the corresponding eigenvalue:

$L_{k}=\sqrt{\lambda_{k}}$	[Eq. 6]

Then, the aspect ratio was defined as:

$\mathsf{Aspect Ratio =}\frac{\mathsf{max(}L_{k})}{\mathsf{min(}L_{k})}$	[Eq. 7]

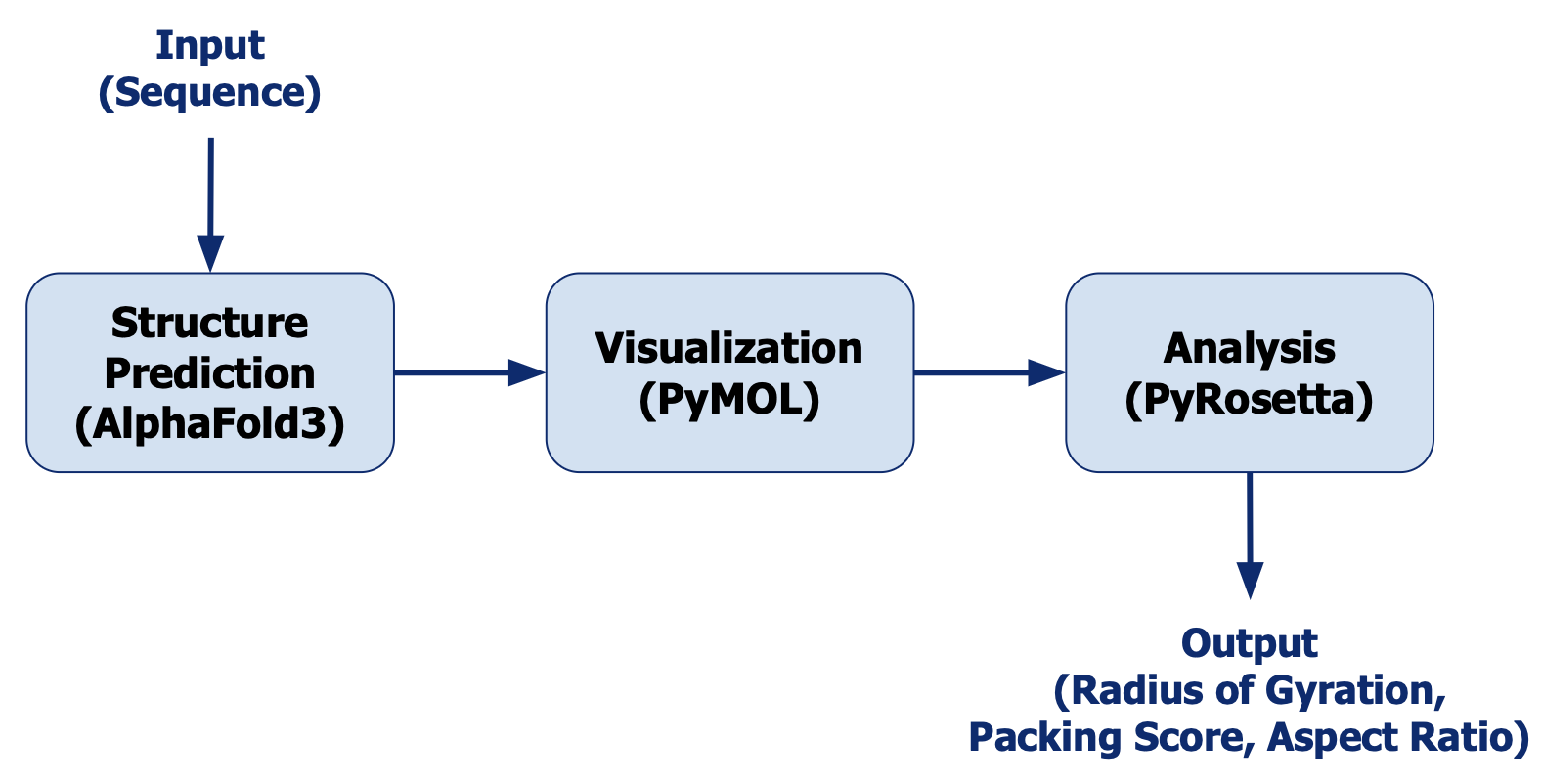

**Figure S1. Computational workflow for AlphaFold3-based analysis of peptide self-assembly.** Peptide sequences and copy numbers were provided as inputs to AlphaFold3 (AF3) to generate predicted supramolecular structures. Predicted structures were subsequently visualized and examined in PyMOL, followed by quantitative structural analysis using PyRosetta to characterize structural and intermolecular features of the predicted assemblies.

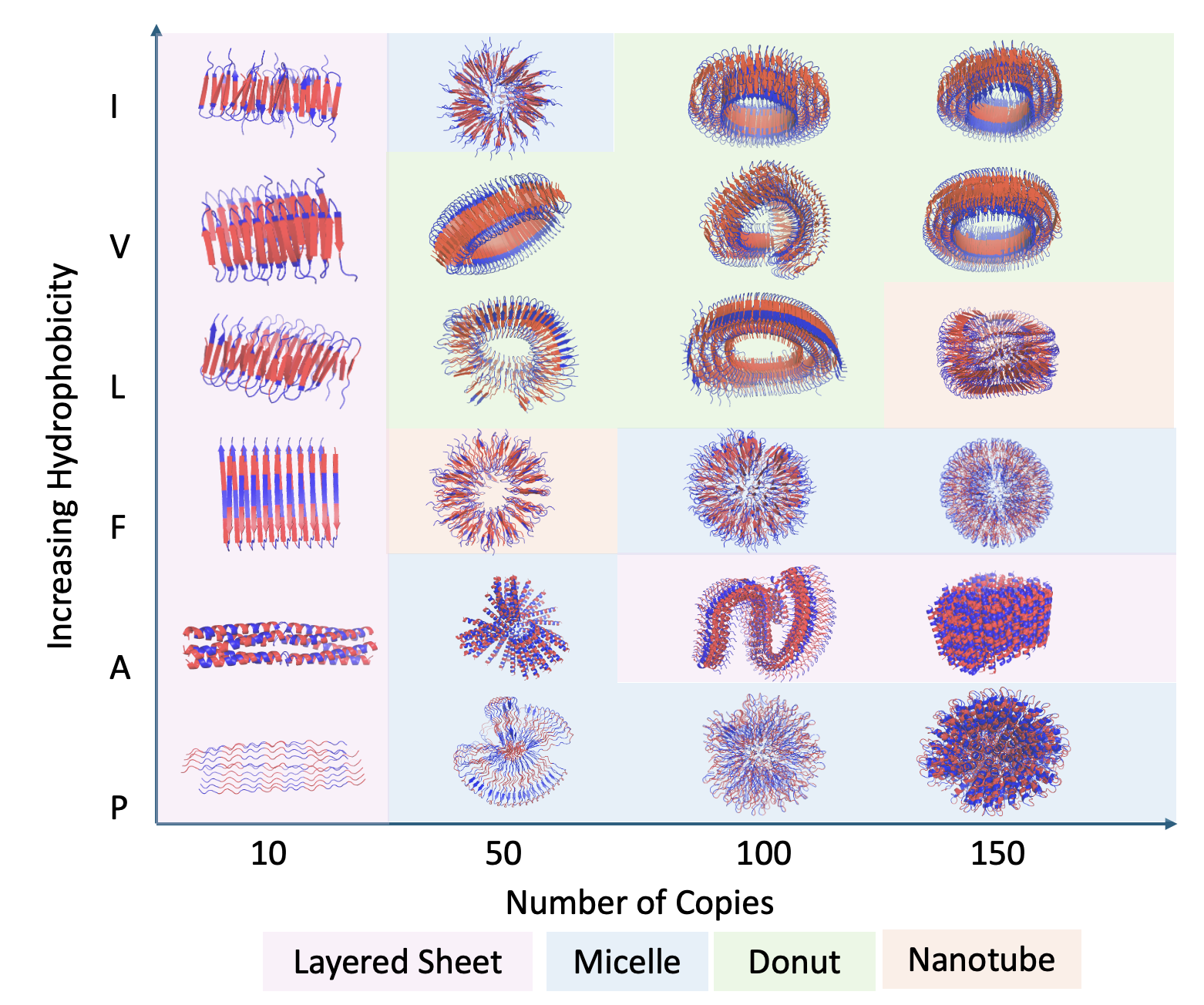

**Figure S2. Phase diagram showing increasing hydrophobicity vs number of copies of (X_4_E_4_)_4_ sequences**, where X represents a hydrophobic amino acid. Beyond fifty copies, we see various micelles, donuts, sheets, and nanotubes. (V_4_E_4_)_4_ formed nano-doughnut structures with hydrophilic edges and hydrophobic cores, suggesting potential tubular growth. Introducing aspartic acid in place of glutamic acid led to flat β-sheet assemblies, possibly forming nanofilaments. Substituting valine with leucine produced hollow vesicles with aqueous interiors. Exchanging aspartic acid for glutamic acid again yielded donut-like structures, but side views revealed cylindrical nanotube formation. (F_4_E_4_)_4_ resembled (L_4_E_4_)_4_ micelles but appeared larger and looser due to the bulkier phenylalanine side chains.

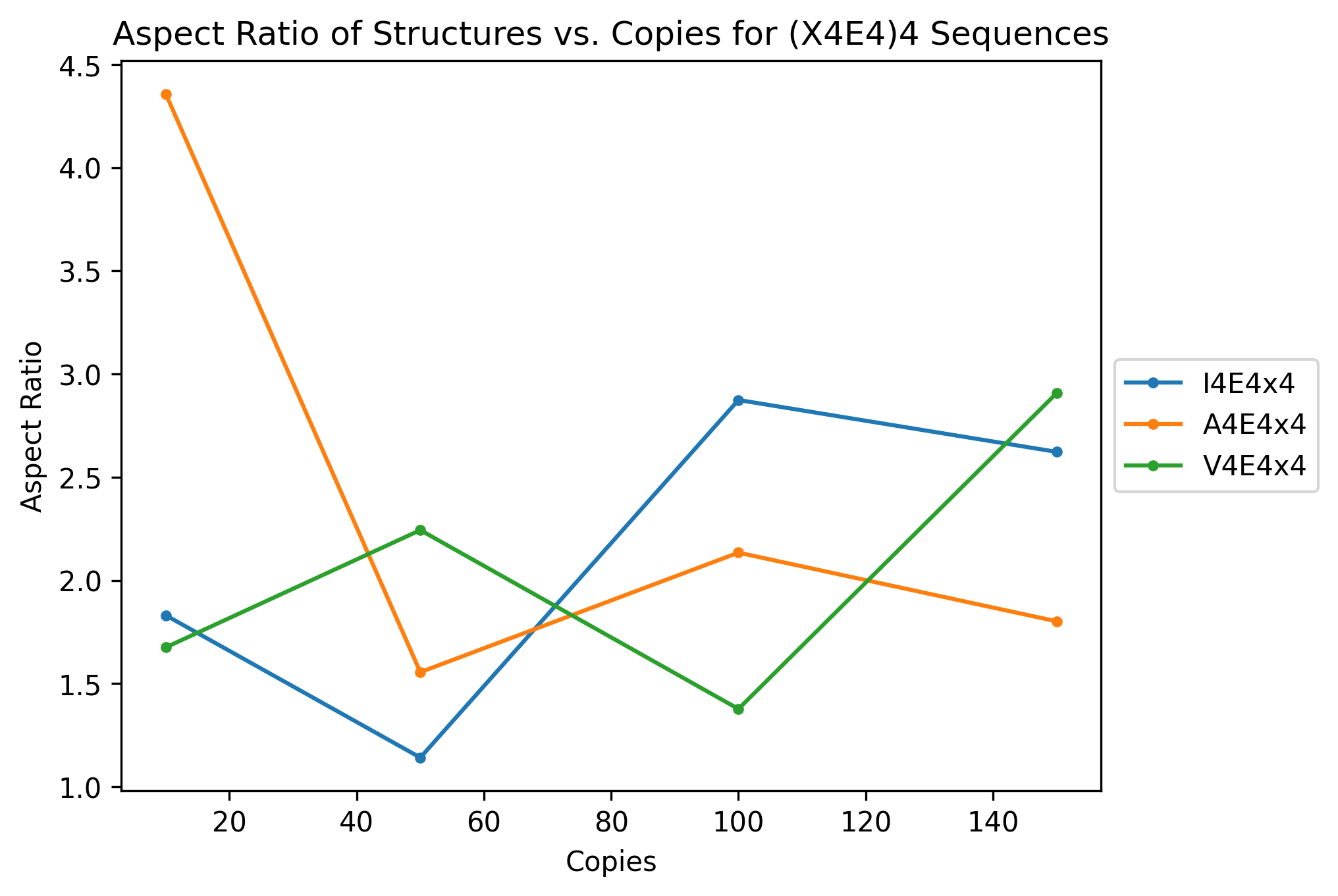

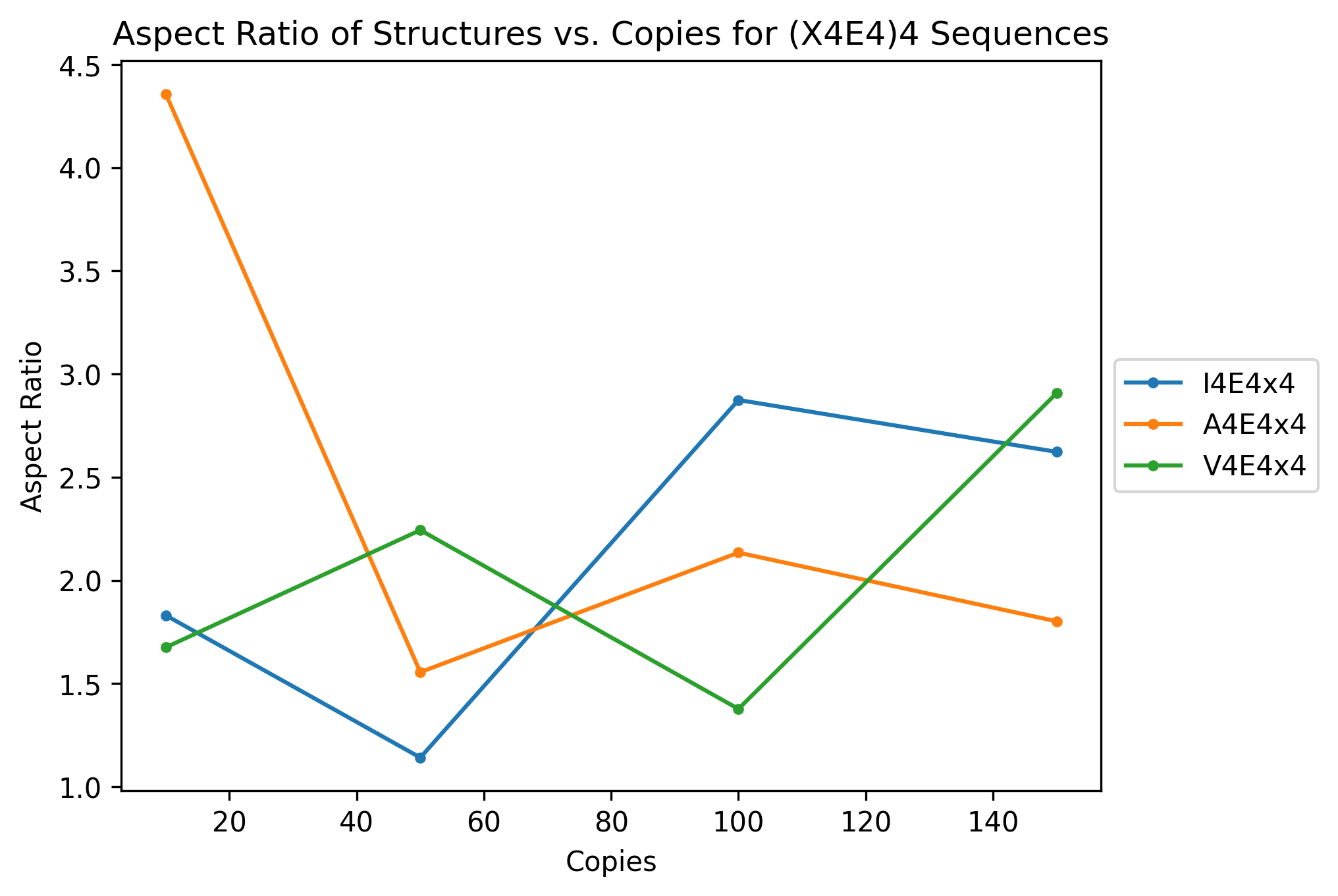

Aspect Ratio

Copies

**Figure S3. Aspect ratio of structures vs copies for (X_4_E_4_)_4_ sequences.** Aspect ratios decreased as copy number increased, indicating a transition from elongated assemblies at low copy numbers toward more compact, isotropic structures.

**(A)	(B)	(C)	(D)**

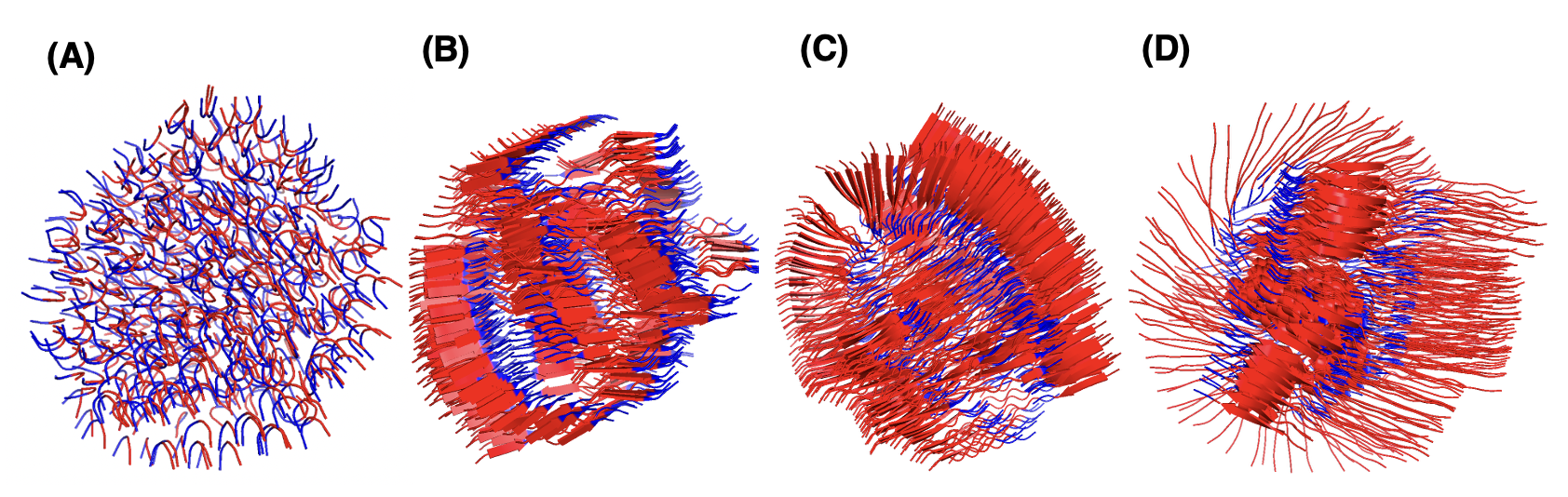

**Figure S4. Adding more valines to the hydrophobic end of the amphiphilic peptide, ranging from 2-8.** (A) V_2_E_2_, 500 copies; (B) V_4_E_2_, 500 copies; (C) V_6_E_2_, 500 copies; (D) V_8_E_2_, 500 copies. As segments lengthened, predicted assemblies showed reduced global order. Overall structures lacked the compartmentalization typical of amphiphilic assemblies.

**(A)	(B)	(C)	(D)**

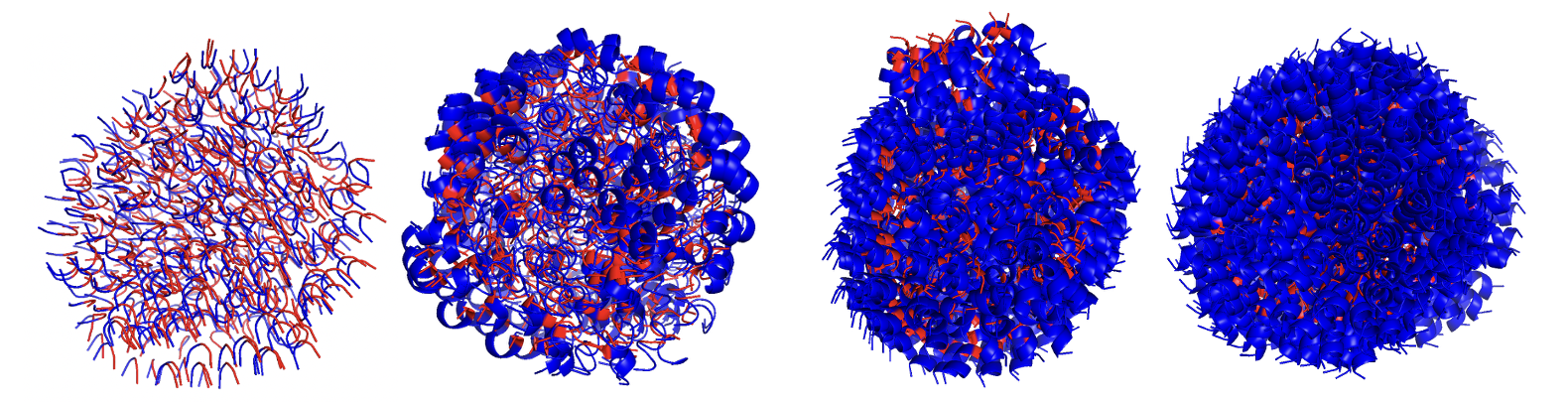

**Figure S5. Adding more glutamines to the hydrophilic end of the amphiphilic peptide, ranging from 2-8.** (A) V_2_E_2_, 500 copies; (B) V_2_E_4_, 500 copies; (C) V_2_E_6_, 500 copies; (D) V_2_E_8_, 500 copies. Increasing hydrophilic residues led to progressively more compact and ordered assemblies.

**(A)	(B)	(C)**

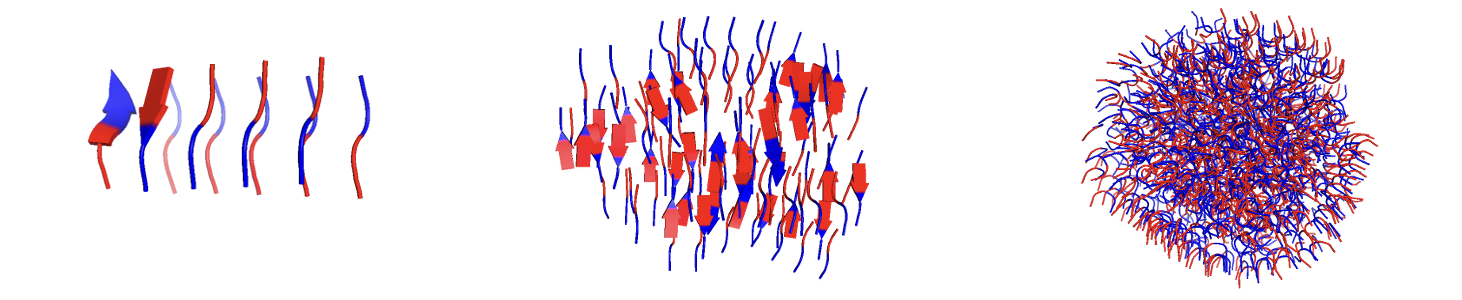

**(D)	(E)	(F)**

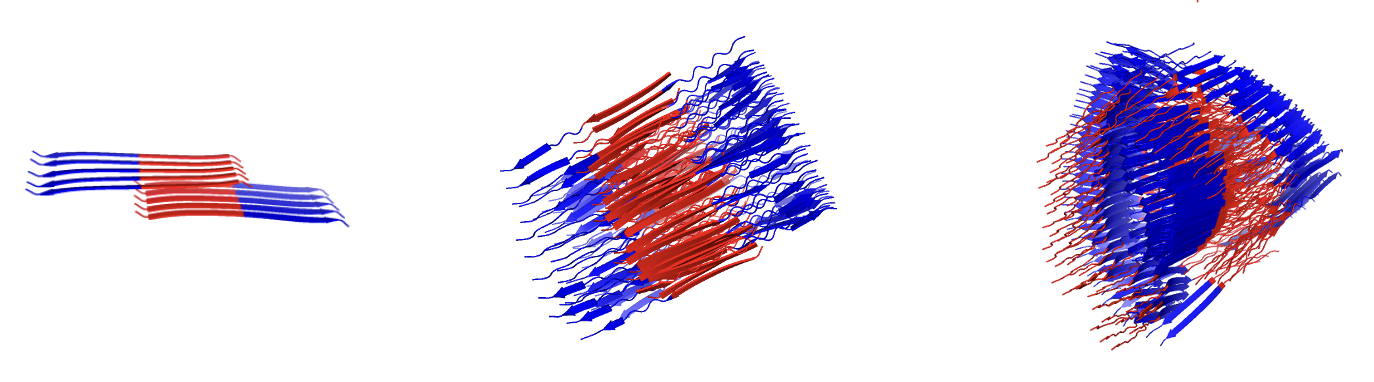

**(G)	(H)	(I)**

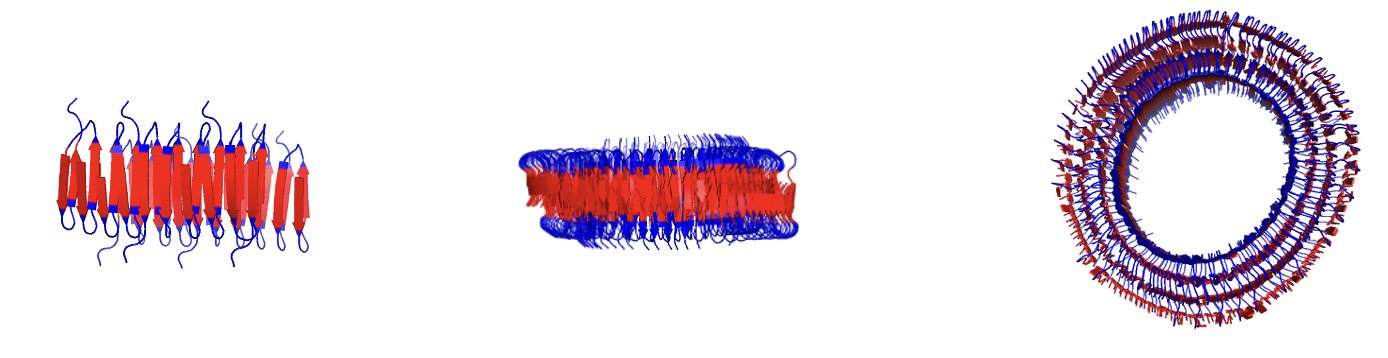

**Figure S6. Examining tetrapeptide, block, and alternating valine and glutamic acid sequences.** (A) V_2_E_2_, 10 copies; (B) V_2_E_2_, 100 copies; (C) V_2_E_2_, 1000 copies; (D) V_9_E_9_, 10 copies; (E) V_9_E_9_, 100 copies; (F) V_9_E_9_, 250 copies; (G) (V_4_E_4_)_4_, 10 copies; (H) (V_4_E_4_)_4_, 100 copies; (I) (V_4_E_4_)_4_, 150 copies. Compared to tetrapeptides, both designs showed improved self-assembly, with hydrophobic cores and hydrophilic shells reflecting the hydrophobic effect. Repeating blocks promoted formation of nano-doughnut structures, contrasting with the sheet-like aggregates of the two-block sequence and the disordered micelles seen with tetrapeptides.

**(A)	(B)	(C)	(D)**

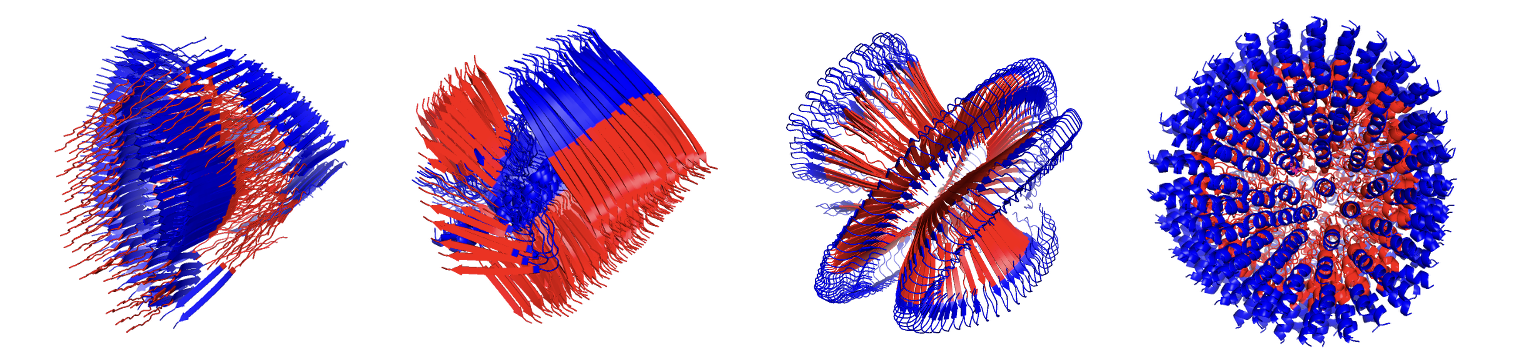

**(E)	(F)	(G)	(H)**

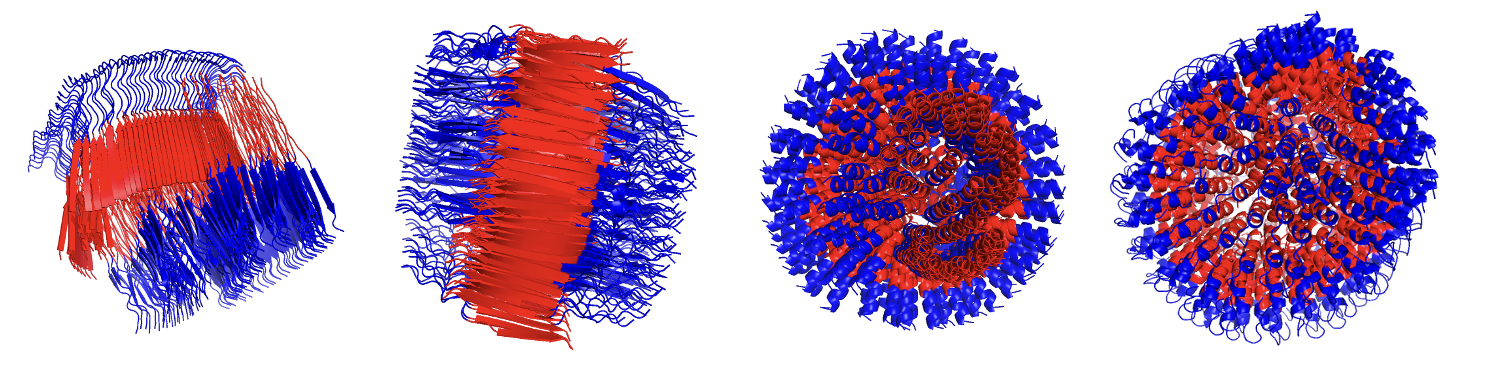

**Figure S7. Effect of input sequence ordering on AlphaFold3-predicted assembly structures using 250 copies of 18-residue peptides with reversed sequence order.** (A) V_9_E_9_; (B) E_9_V_9_;(C) F_9_E_9_; (D) E_9_F_9_; (E) I_9_E_9_; (F) E_9_I_9_; (G) L_9_E_9_; (H) E_9_L_9_. Notably, F_9_E_9_ formed β-sheet micelles, while E_9_F_9_ formed α-helical micelles, suggesting sequence order influences assembly. Most inversions maintained hydrophobic cores and hydrophilic exteriors. Sequence inversion shifted the spatial distribution of key intermolecular interactions, disrupting cooperative packing and impacting the formation of a thermodynamically stable core.

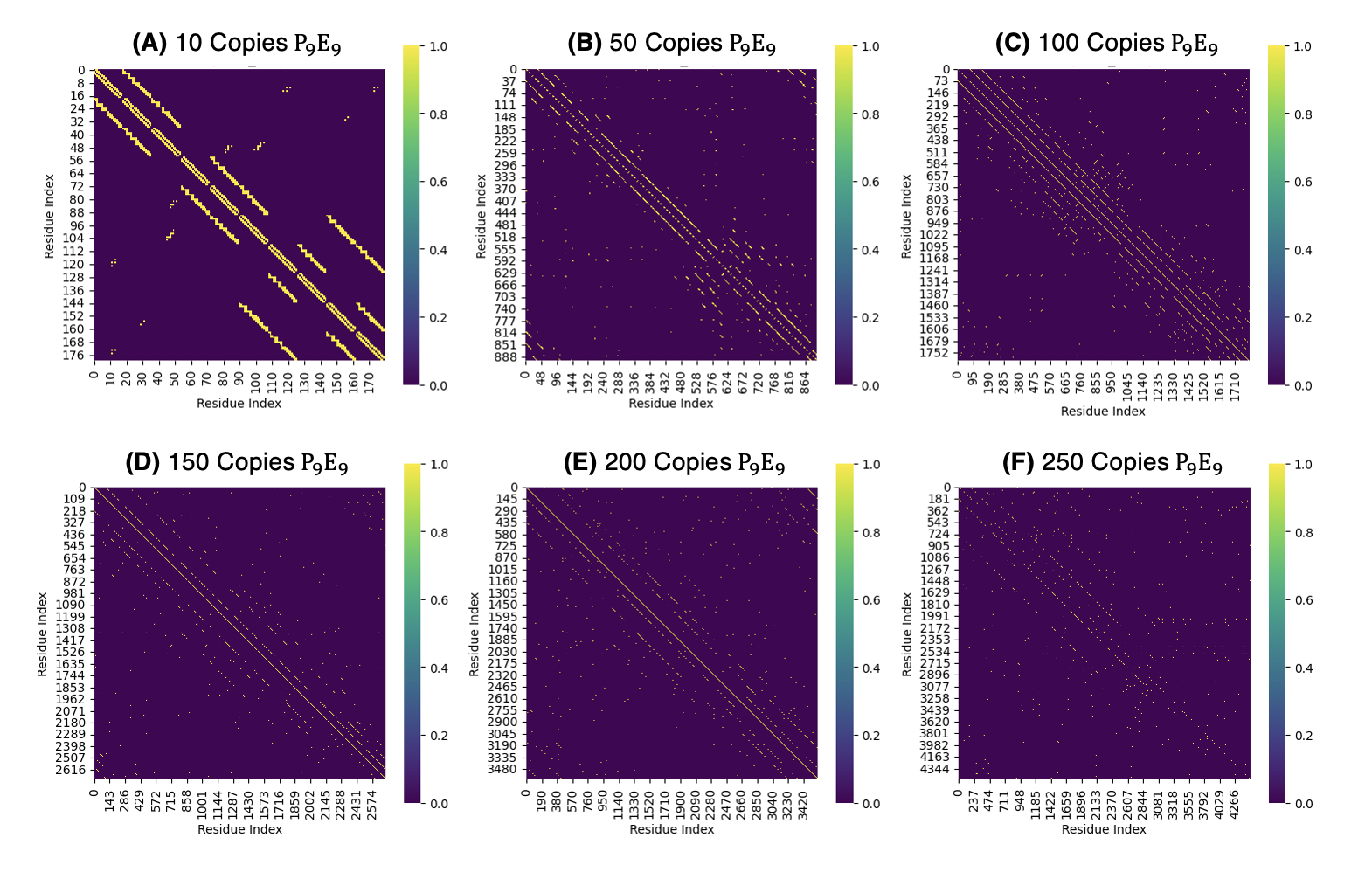

**Figure S8. Contact maps of P_9_E_9_ assemblies predicted by AlphaFold3 across varying peptide copy numbers.** (A) 10 copies; (B) 50 copies; (C) 100 copies; (D) 150 copies; (E) 200 copies; (F) 250 copies. Contact maps are plotted as a function of residue index, with the diagonal bands representing recurring contacts within and between peptide chains. At lower copy numbers, the assemblies exhibit strong, well-defined diagonal contact patterns, while increasing copy number results in a more diffuse distribution of additional intermolecular contacts. Copy number represents the number of peptide chains specified in the AlphaFold3 prediction and should not be interpreted as a direct measure of bulk concentration.

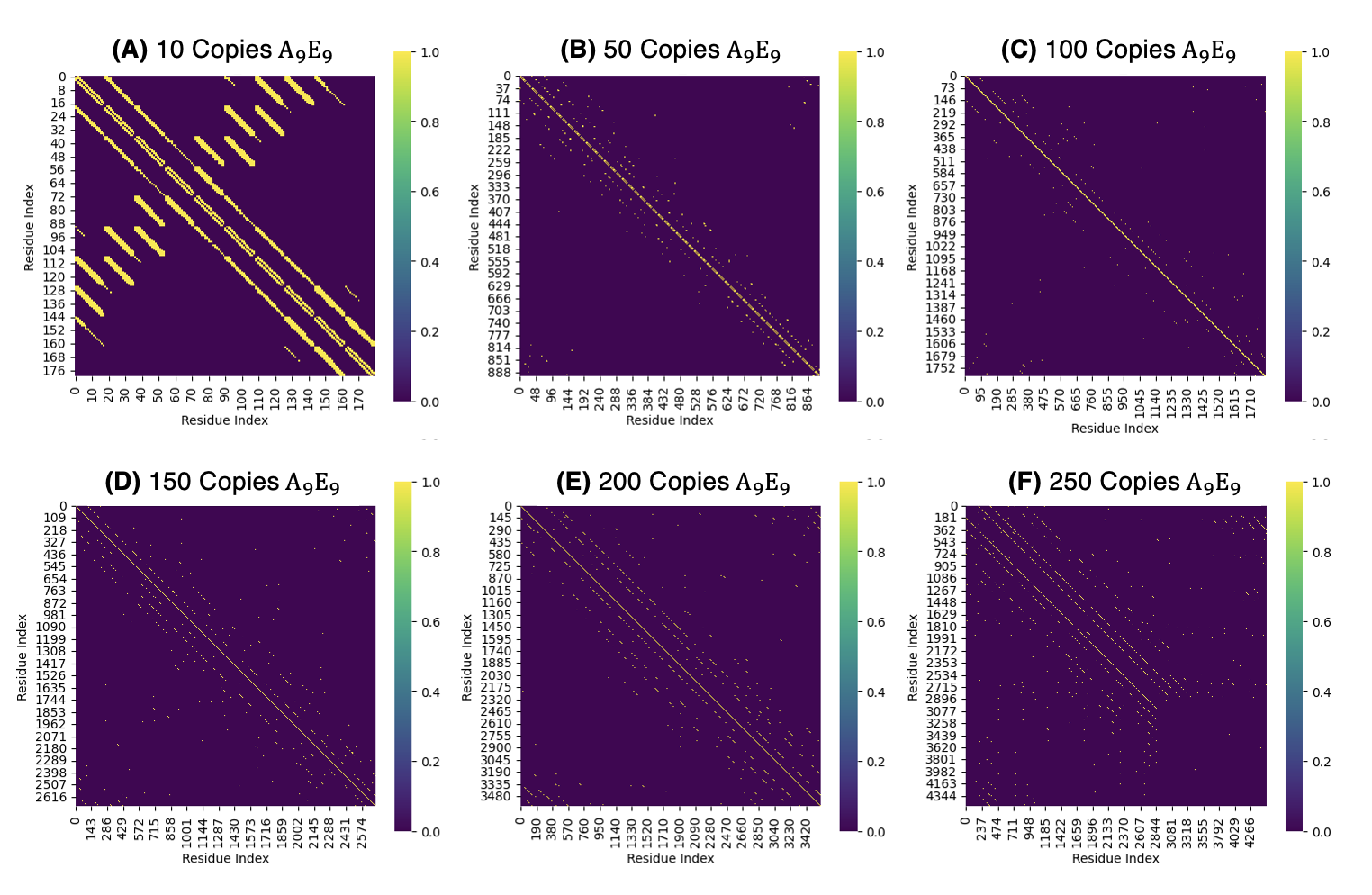

**Figure S9. Contact maps of A_9_E_9_ assemblies predicted by AlphaFold3 across varying peptide copy numbers.** (A) 10 copies; (B) 50 copies; (C) 100 copies; (D) 150 copies; (E) 200 copies; (F) 250 copies. Contact maps are plotted as a function of residue index, with the diagonal bands representing recurring contacts within and between peptide chains. At 10 copies, A_9_E_9_ shows extensive, highly structured contact bands, indicating a more ordered packing arrangement. As copy number increases, the contact patterns become progressively more dispersed, with fewer continuous high-intensity bands and more isolated intermolecular contacts. These changes suggest that higher-copy assemblies sample a broader range of packing interactions while retaining a persistent diagonal signature associated with recurring local organization. Copy number refers only to the number of peptide chains used in the AlphaFold3 prediction and does not correspond directly to solution concentration.

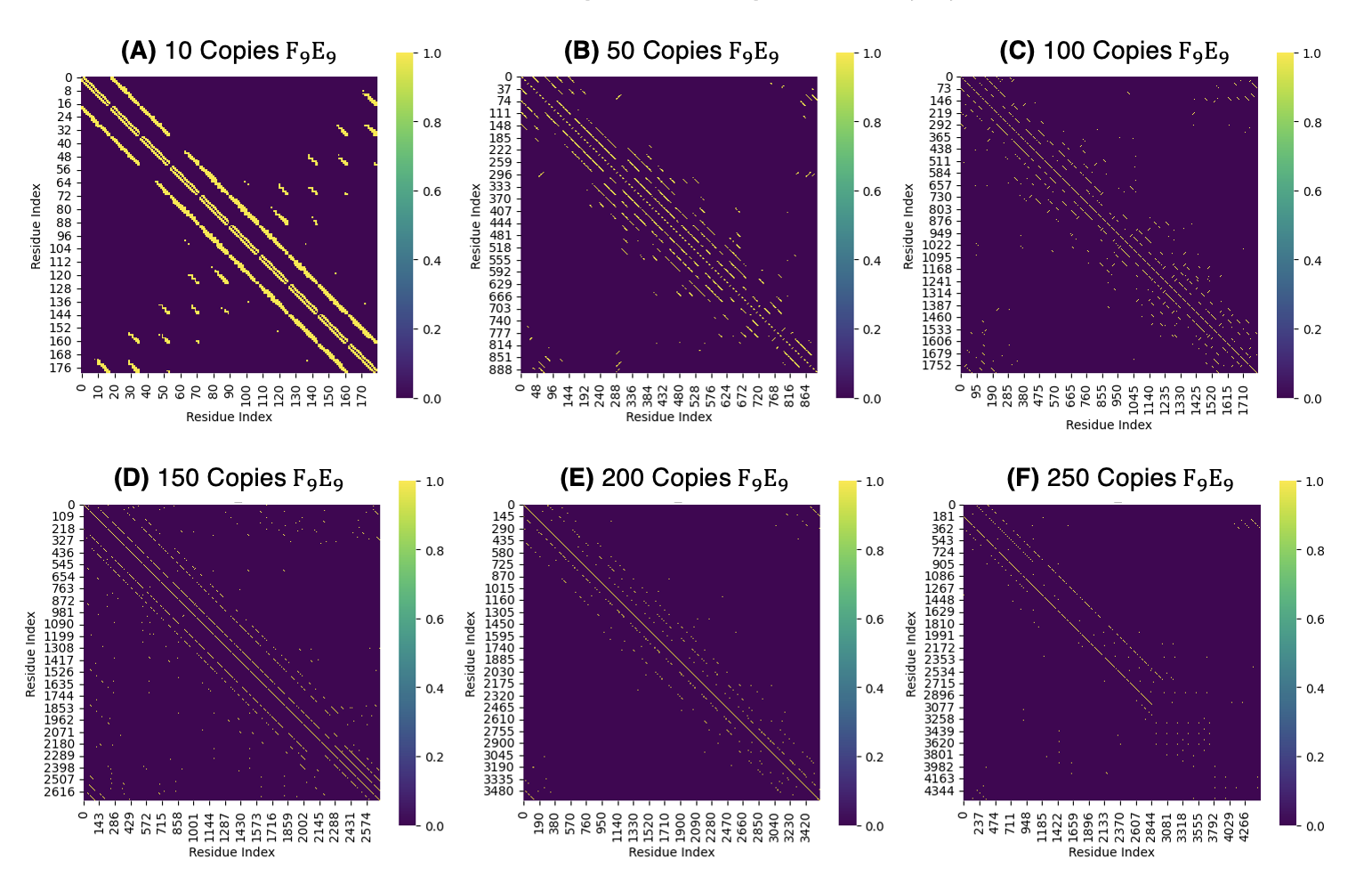

**Figure S10. Contact maps of F_9_E_9_ assemblies predicted by AlphaFold3 across varying peptide copy numbers.** (A) 10 copies; (B) 50 copies; (C) 100 copies; (D) 150 copies; (E) 200 copies; (F) 250 copies. Contact maps are plotted as a function of residue index, with the diagonal bands representing recurring contacts within and between peptide chains. Increasing copy number produces progressively more distributed and less densely defined intermolecular contacts, with the strongest contact patterns becoming concentrated along the diagonal. These results demonstrate that increasing peptide copy number alters the predicted contact network and packing arrangement of the F₉E₉ assembly; however, copy number should be interpreted as a model input rather than a direct measure of bulk concentration.

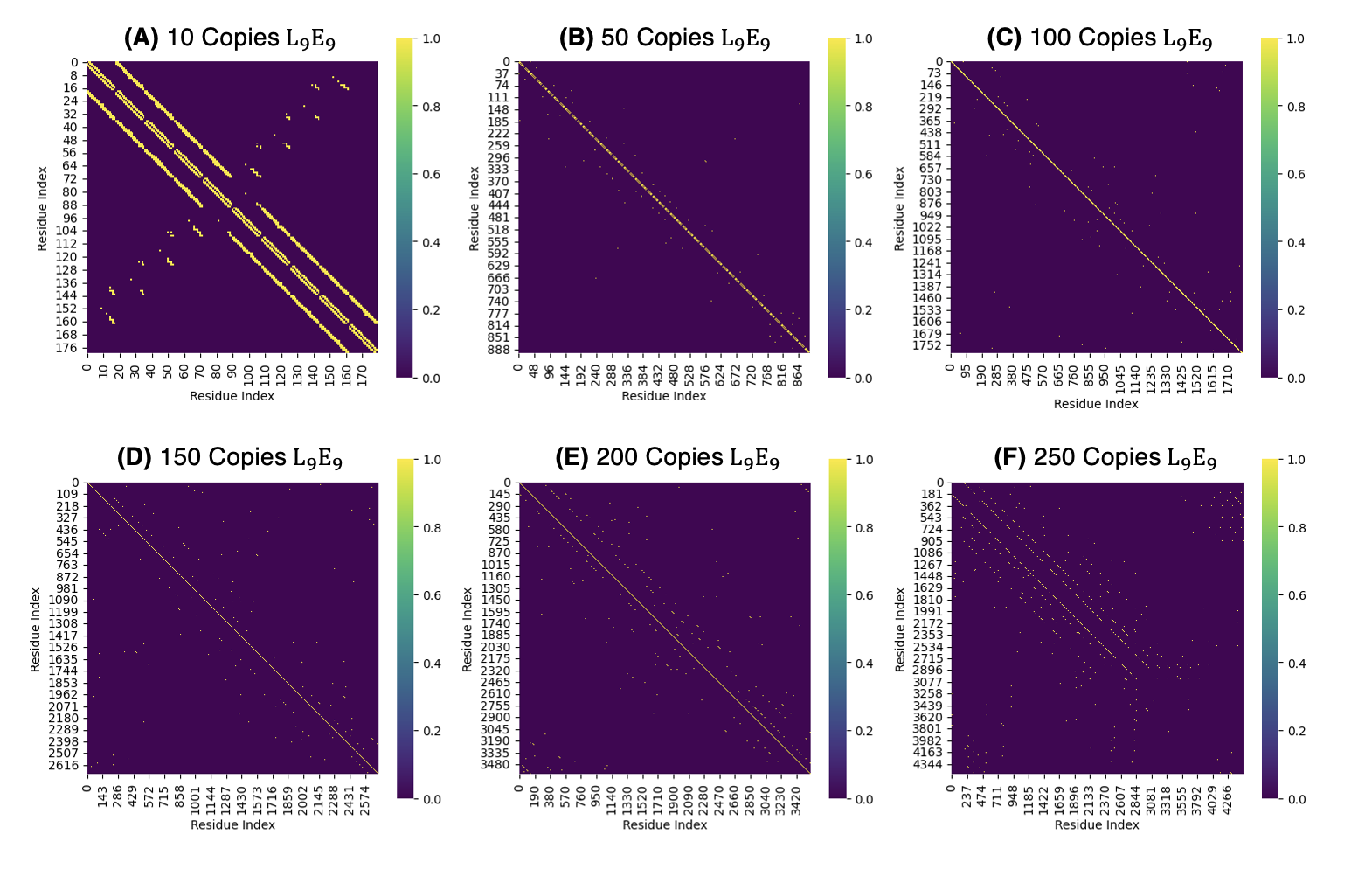

**Figure S11. Contact maps of L_9_E_9_ assemblies predicted by AlphaFold3 across varying peptide copy numbers.** (A) 10 copies; (B) 50 copies; (C) 100 copies; (D) 150 copies; (E) 200 copies; (F) 250 copies. Contact maps are plotted as a function of residue index, with the diagonal bands representing recurring contacts within and between peptide chains. At 10 copies, L₉E₉ exhibits several strong diagonal contact bands and additional intermolecular contacts. With increasing copy number, these additional contacts become less prominent, while the primary diagonal contact pattern persists. At 250 copies, a more distributed network of intermolecular contacts is observed. These results demonstrate changes in the predicted contact network with increasing peptide copy number; however, copy number represents the number of peptide chains specified in the AlphaFold3 prediction and should not be interpreted as a direct measure of bulk concentration.

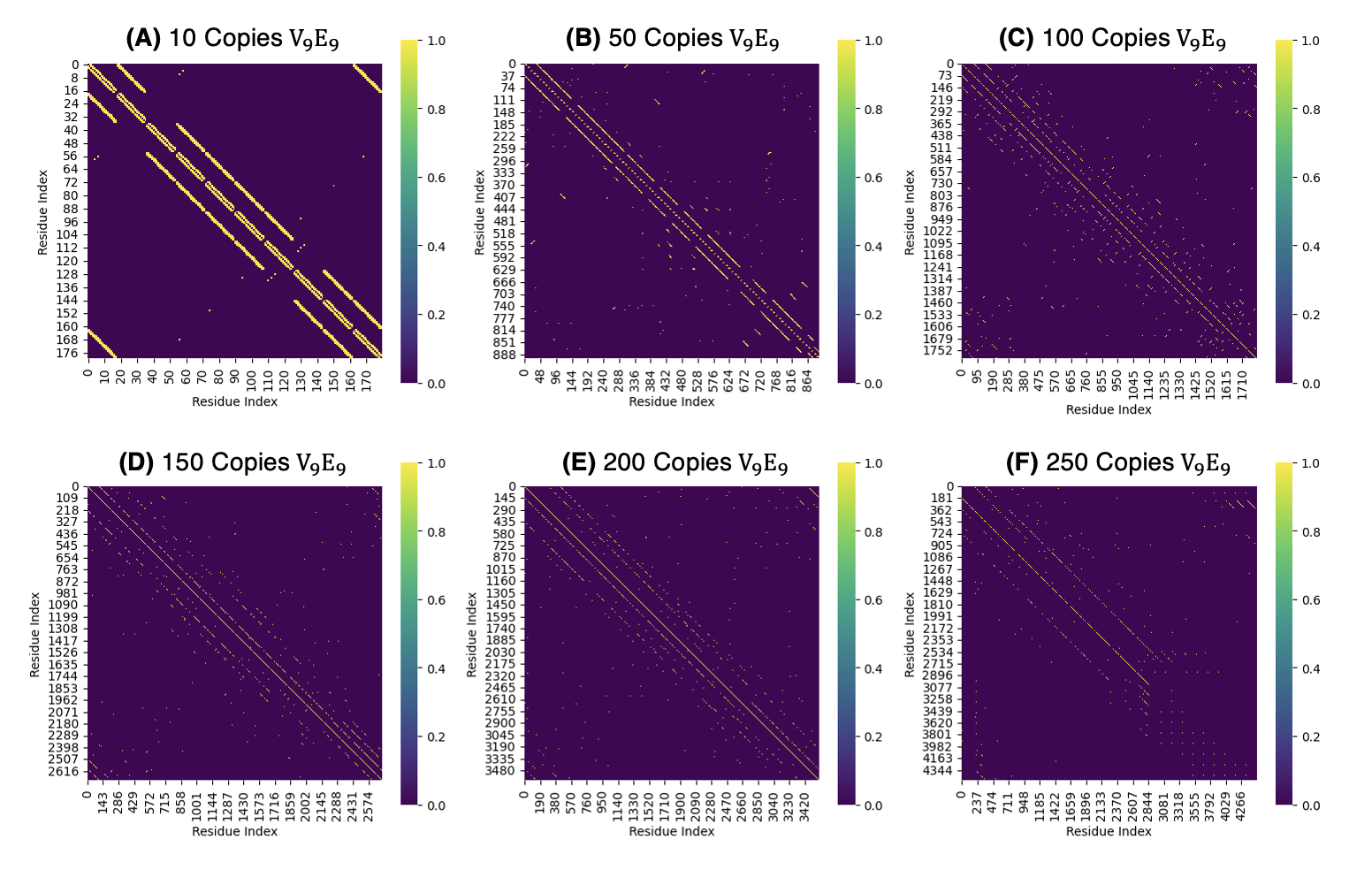

**Figure S12. Contact maps of V_9_E_9_ assemblies predicted by AlphaFold3 across varying peptide copy numbers.** (A) 10 copies; (B) 50 copies; (C) 100 copies; (D) 150 copies; (E) 200 copies; (F) 250 copies. Contact maps are plotted as a function of residue index, with the diagonal bands representing recurring contacts within and between peptide chains. At lower copy numbers, the assemblies display strong, well-defined diagonal patterns, whereas increasing copy number produces a more diffuse distribution of intermolecular contacts. The persistence of diagonal bands across copy numbers suggests recurrent local packing interactions within the predicted assemblies. Copy number refers to the number of peptide chains specified in the AlphaFold3 prediction and should not be interpreted as a direct measure of bulk concentration.

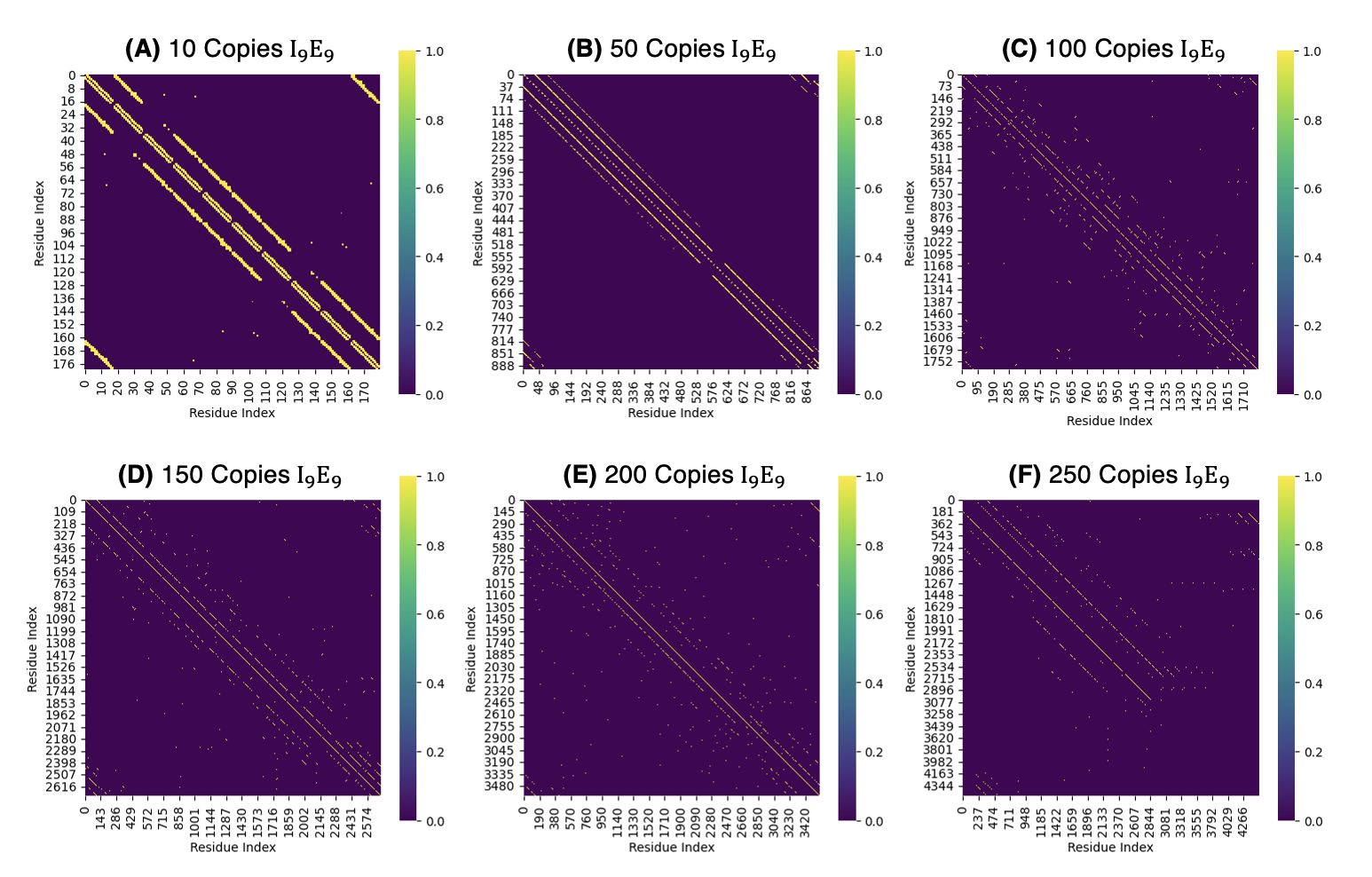

**Figure S13. Contact maps of I_9_E_9_ assemblies predicted by AlphaFold3 across varying peptide copy numbers.** (A) 10 copies; (B) 50 copies; (C) 100 copies; (D) 150 copies; (E) 200 copies; (F) 250 copies. Contact maps are plotted as a function of residue index, with the diagonal bands representing recurring contacts within and between peptide chains. At lower copy numbers, the assemblies exhibit strong, well-defined diagonal contact patterns, while increasing copy number results in a more diffuse distribution of additional intermolecular contacts. The persistence of diagonal bands across copy numbers suggests recurring local packing interactions within the predicted assemblies. Copy number represents the number of peptide chains specified in the AlphaFold3 prediction and should not be interpreted as a direct measure of bulk concentration.
